# Serotonergic modulation of walking in *Drosophila*

**DOI:** 10.1101/753624

**Authors:** Clare E. Howard, Chin-Lin Chen, Tanya Tabachnik, Rick Hormigo, Pavan Ramdya, Richard S. Mann

## Abstract

To navigate complex environments, animals must generate highly robust, yet flexible, locomotor behaviors. For example, walking speed must be tailored to the needs of a particular environment: Not only must animals choose the correct speed and gait, they must also rapidly adapt to changing conditions, and respond to sudden and surprising new stimuli. Neuromodulators, particularly the small biogenic amine neurotransmitters, allow motor circuits to rapidly alter their output by changing their functional connectivity. Here we show that the serotonergic system in the vinegar fly, *Drosophilamelanogaster*, can modulate walking speed in a variety of contexts and in response to sudden changes in the environment. These multifaceted roles of serotonin in locomotion are differentially mediated by a family of serotonergic receptors with distinct activities and expression patterns.

## Introduction

Insects have a remarkable capacity to adapt their locomotor behaviors across a wide range of environmental contexts and to confront numerous challenges. For example, they can walk forwards, backwards, and upside down, navigate complex terrains, and rapidly recover after injury [1-9]. To achieve this wide range of behaviors, insects regulate their global walking speed and kinematic parameters, allowing them to modify stereotyped gaits as needed [3,5-7,9,10]. Because overlapping sets of motor neurons and muscles are recruited for all of these behaviors, animals must be able to rapidly modulate the circuit dynamics that control locomotor parameters [11-13] (reviewed in [14]).

As with limbed vertebrates, most insects use multi-jointed legs to walk [8,10,15-17]. Locomotor circuits that orchestrate these complex gaits are located in the ventral nerve cord (VNC), a functional analogue of the vertebrate spinal cord that includes three pairs of thoracic neuromeres (T1, T2, and T3) that coordinate the movements of the three pairs of thoracic legs [1-9,18-20]. The insect VNC receives descending commands from the brain and sends motor output instructions via motor neurons to peripheral musculature [3,5-7,9,10,19]. Leg motor neuron dendrites innervate the leg neuropils within the VNC and their axons exit the VNC to synapse onto muscles in the appendages [11-13,21,22]. Sensory neurons, which convey proprioceptive and tactile information, project axons from the appendages to the VNC by these same fiber tracts, where they arborize in the leg neuropils [14,23,24] (Figure 1A). Notably, the VNC is capable of executing coordinated leg motor behaviors, such as walking and grooming, even in decapitated animals [8,10,15-17,25]. Thus, the VNC harbors neural networks that can drive the coordinated flexion and extension of each leg joint and, therefore, walking gaits [8,10,17,20,26].

**Figure 1.**
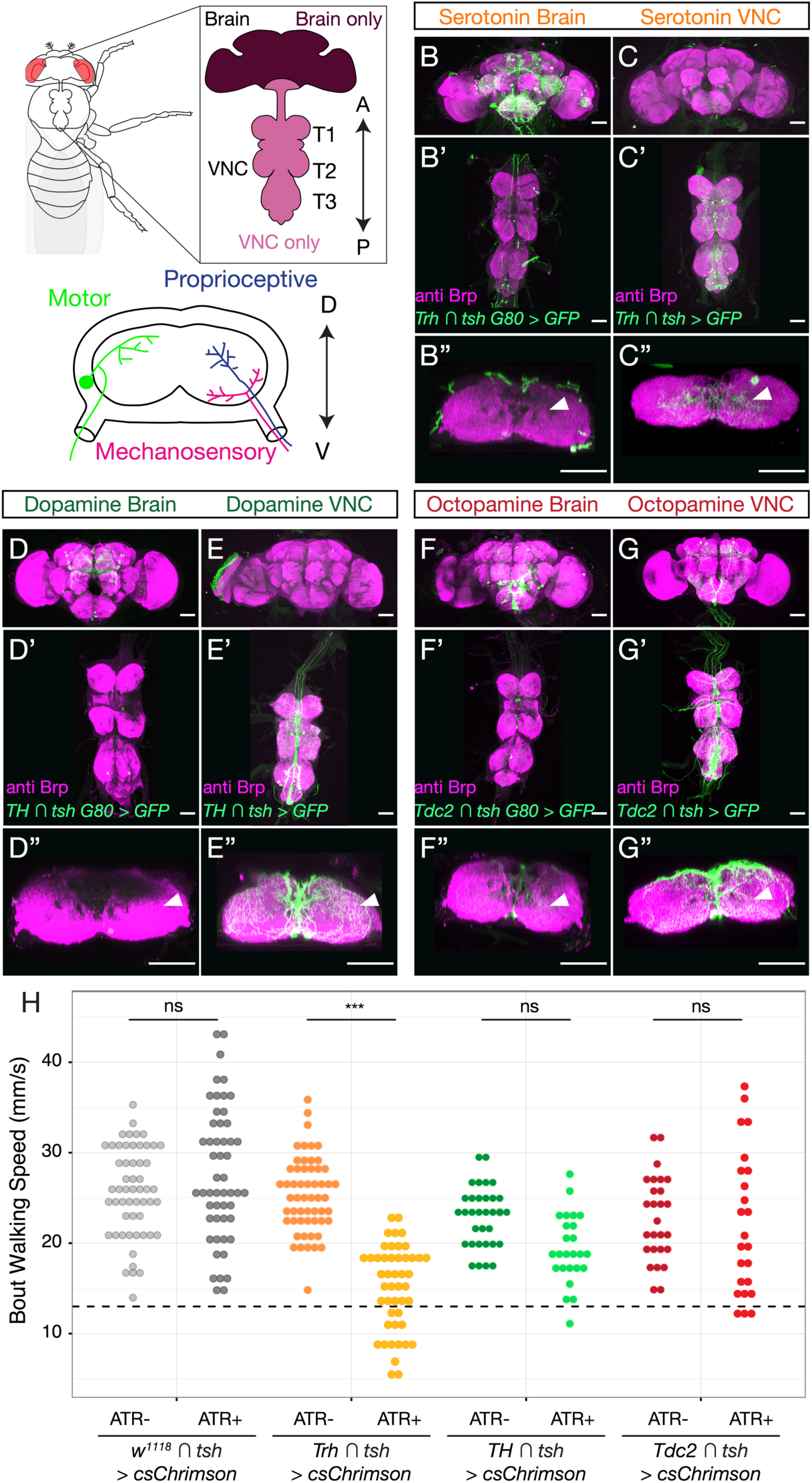
Neuromodulators in the fly CNS. **A**. The adult *Drosophila* CNS is composed of the brain and the VNC, which consists of three thoracic neuropils (T1, T2 and T3), each of which corresponds to a pair of adult legs, and an abdominal ganglion. Anterior-posterior axis specified. *Lowerpanel:* Each thoracic neuropil contains the projections of locomotor circuit components, including motor neurons that send axons to leg muscles and sensory neurons that convey mechanosensory and proprioceptive information from the legs (schematized here in cross section). Dorsal – ventral axis specified. **B-G**. Maximum intensity projections show the expression patterns driven by Gal4 lines labeling either brain-derived (B, D, F, *Gal4* intersected with *tshGal80*) or VNC-derived (C, E, G, *Gal4* intersected with *tsh*) serotonergic (B-C, *TrhGal4*), dopaminergic (D-E, *THGal4*), or octopaminergic/tyraminergic (F-G, *Tdc2Gal4*) neurons. (B’’-G’’) Projection of a subset of cross sections of the VNC shows innervation of the T1 neuropil. All scale bars are 50 um. **H**. Optogenetic activation of serotonergic (*Trh-Gal4*) neurons in the *Drosophila* VNC, but not dopaminergic (*TH-Gal4*) or octopaminergic/tyraminergic (*Tdc2-Gal4*) neurons, slows walking speed compared to all-trans-retinal (ATR) negative and non-Gal4 (*w*^*1118*^) controls. These activation experiments were carried out using the Flywalker assay (Mendes et al., 2013; see Figure S3A for a schematic). Statistics computed using a Kruskal-Wallis test with the Dunn-Sidak correction for multiple comparisons * <.05 **<.01 ***<.001. N walking bouts (animals) *w*^*1118*^ ATR-55 (14-31); *w*^*1118*^ ATR+ 52 (14-36); *Trh* ATR-56 (12-30); *Trh* ATR+ 47 (10-23); *TH* ATR-33 (10-24); *TH* ATR+ 25 (10-26); *Tdc2* ATR-27 (10-27); *Tdc2* ATR+ 24 (10-25).

Numerous studies have shown that sensory input from the legs are required for robust and stereotyped locomotor patterns, regulating both the timing and magnitude of locomotor activity and also facilitating coordination between legs [6,8,10,12,27-29]. However, sensory feedback cannot be the only means for tuning locomotion: mutation of proprioceptive receptors or even deafferenting limbs does not block coordinated walking [10,15,18,30-33]. Beyond sensory feedback-driven tuning of gait patterns, larger behavioral changes must be accomplished by other circuits. These likely include neuromodulatory systems, including the monoamines dopamine, norepinephrine, and serotonin, which are highly conserved throughout the animal kingdom.

Monoamines have been shown to modulate, and even induce, the activities of central pattern generating (CPG) motor circuits. In crustaceans, for example, neuromodulation causes the gastric CPG to generate distinct rhythmic activity patterns from the same neural network to address distinct behavioral demands [34-39] (reviewed in [13,40]). Remarkably, the same neuromodulatory systems appear to play similar roles across species. For example, serotonin has been shown to slow locomotor rhythms in animals as diverse as the lamprey, cat, and locust [41-43]. In the vinegar fly, *Drosophilamelanogaster*, monoamine neurotransmitters have also been shown to modulate walking behavior. In addition to slowing walking speed, serotonin modulates sleep and anxiety-related motor behaviors [3,5,6,9,10,44-48]. Dopamine, in contrast, has been linked to hyperactivity [25,49-52]. Octopamine has been shown to mediate starvation induced hyperactivity, and in its absence animals walk more slowly [8,53,54]. As each of these neuromodulatory systems plays a variety of roles in regulating complex behaviors, it has thus far been challenging to tease apart which of the effects on walking behavior are due to direct modulation of motor circuitry or are a secondary consequence of modulating higher order circuits in the brain.

In this work, we show that the serotonergic neurons within the VNC can modulate walking speed in a context-independent manner as well as in response to startling stimuli. Additionally, we demonstrate that these modulatory effects are enacted through serotonin’s action via specific receptors that are expressed in different parts of the locomotor circuit. Together, these findings reveal that neuromodulatory systems regulate multiple aspects of complex behaviors such as walking at multiple time scales, allowing animals to effectively respond to rapidly changing environments.

## Results

### VNC serotonergic neurons arborize within the leg neuropils

To identify neuromodulatory neurons that might play a role in modulating walking behavior we drove expression of a fluorescent reporter with Gal4 under the control of promoters encoding key synthetic enzymes for each neuromodulatory system – *Tryptophan hydroxylase* (*Trh* for serotonin (5-HT) [55]); *tyrosinehydroxylase* (*TH* or *ple* (*pale*) for dopamine [56]); and *Tyrosine decarboxylase2* (*Tdc2* for octopamine and tyramine [57]). All of these drivers show extensive expression in cells both within the VNC and the brain, with processes that densely innervate VNC leg neuromeres (Figure 1) [55].

To determine whether local VNC neurons or descending neurons originating in the brain innervate the leg neuropils, we used genetic intersectional tools to limit the expression of these Gal4 lines to either the brain or VNC (Figure 1A). These experiments show that neuromodulatory innervation of the leg neuropils arises almost entirely from VNC interneurons and not from descending neurons in the brain (Figure 1B-G). Moreover, in many cases, these VNC neurons extensively innervate the leg neuropils. Thus, VNC neuromodulatory neurons are well-positioned to directly modulate VNC locomotor circuits. These intersectional genetic tools therefore allow us to assess the behavioral role of local VNC neuromodulation independently of modulation within higher brain regions.

### Activation of VNC serotonergic neurons slows walking speed

Previous studies showed that neuromodulatory systems can regulate walking [8,25,47,49,51,54,58], but could not dissociate the relative contribution of brain and VNC neuromodulatory subpopulations. Here we addressed whether neuromodulatory neurons in the VNC alone are sufficient to modulate walking behavior. We specifically optogenetically activated these neurons and found that activation of serotonergic VNC populations, but not dopaminergic or octopaminergic/tyraminergic VNC subpopulations, altered the average speed at which animals walk (Figure 1H).

Based on these results, we focused the remainder of our analysis on VNC serotonergic neurons (5-HT^VNC^). To validate the fidelity of our serotonergic Gal4 driver line, and to rule out co-secretion of other neurotransmitters, we performed immunostaining for markers of serotoninergic (5-HT), dopaminergic (TH), octopaminergic/tyraminergic (Tdc2), glutamatergic (VGlut), cholinergic (ChAT), and GABAergic (GABA) neurons (Figure S1). These experiments demonstrate that the *Trh-Gal4* line drives expression in 5-HT-expressing neurons, and that these neurons do not express any of the other neurotransmitters we surveyed, suggesting that they are primarily serotonergic.

We next confirmed the effects of activating 5-HT^VNC^ neurons by studying animals freely walking within an arena. This allowed us to measure not only an animal’s speed, but also its walking frequency, angular velocity, and preferred position within the arena (Figure S2A-C). As with our initial experiments, activation of 5-HT^VNC^ neurons is sufficient to produce a rapid slowing of average walking speed in this paradigm (Figure 2A). Interestingly, activation of 5-HT^VNC^ neurons does not change the overall amount of time animals spend walking, suggesting that speed changes are not simply due to a decrease in overall activity, but instead reveal a bias towards slower walking speeds (Figure S2D and F). Unlike a previous study showing that overexpressing the serotonin transporter in all neurons caused flies to move away from the edge of the arena [46], we see no effect on the distribution of animals within the arena when we limit the activation to 5-HT^VNC^ neurons (Figure S2D). We also find that activation of 5-HT^VNC^ neurons decreases the absolute angular velocity of walking flies (Figure S2D). Thus, although these flies walk slower, they also walk straighter than control flies. This latter observation is unexpected, because straighter trajectories are usually correlated with faster walking speeds (Figure S2F) [10].

**Figure 2.**
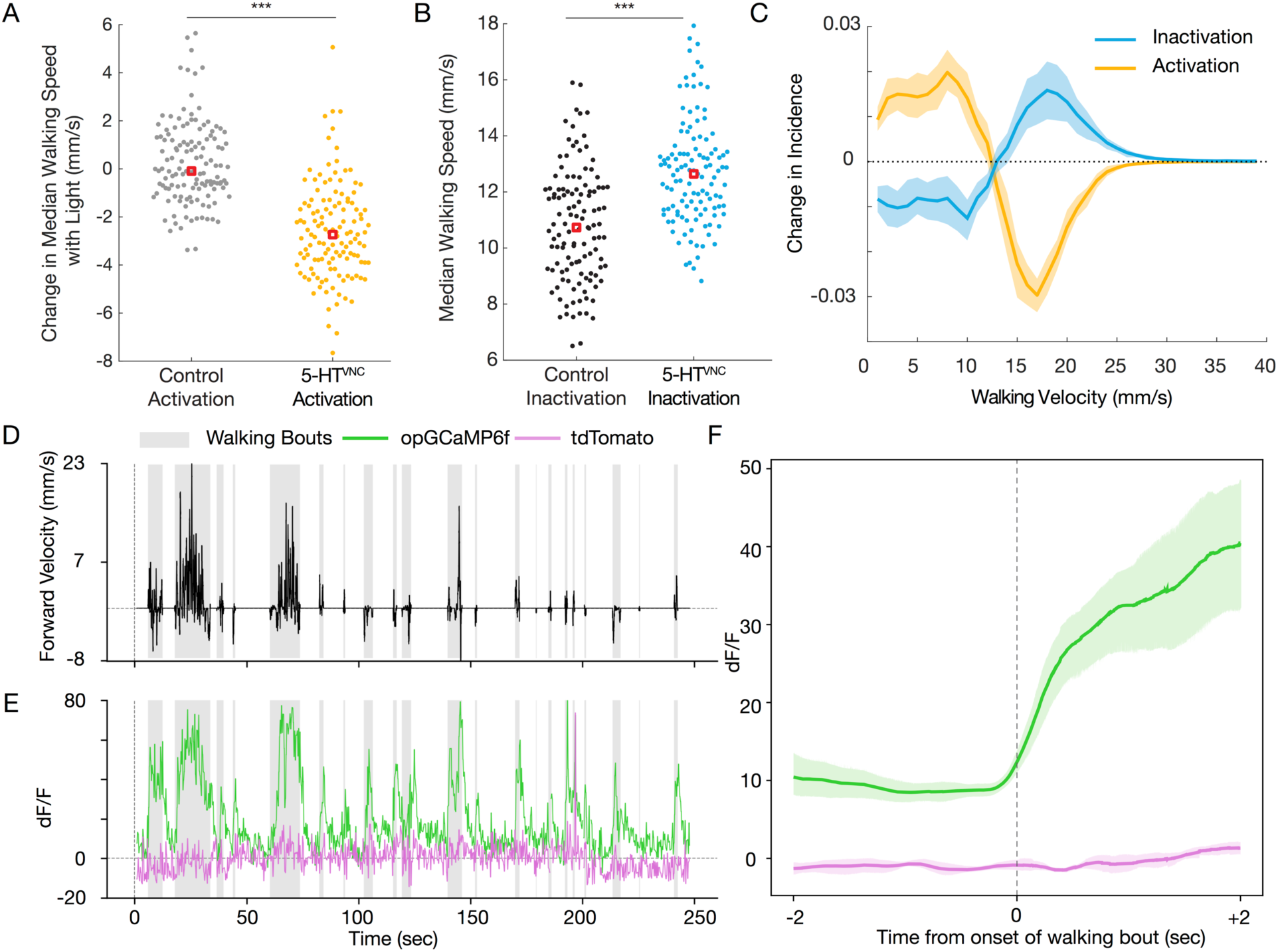
5-HT^VNC^ neurons modulate walking speed. **A**. Activation of 5-HT^VNC^ neurons (*Trh⋂tsh>csChrimson* fed with ATR) causes animals to walk slower than background matched non-Gal4 controls (*w*^*1118*^ *⋂tsh>csChrimson* fed with ATR). Both genotypes were fed all-trans-retinal. Genotypes were compared using Kruskal-Wallis test, ***p<.001. N = 130 animals for each condition. **B**. Inactivation of 5-HT^VNC^ neurons (*Trh⋂tsh>Kir2*.*1*) causes animals to walk faster than genetically matched non-Gal4 controls (*w*^*1118*^ *⋂tsh>Kir2*.*1*). Genotypes were compared using a Kruskal-Wallis test, ***p<.001. N=119 animals per genotype. **C**. The distribution of velocity shifts caused by activation and inhibition of 5-HT^VNC^ neurons are symmetrical. Differences in population average histograms were calculated between control and experimental genotypes and were fit with 95% confidence intervals via bootstrapping. For activation experiments, behavior of *w*^*1118*^ *⋂tsh>csChrimson* flies fed with ATR was compared to that of *Trh⋂tsh>csChrimson* flies also fed with ATR for the light on period only. **D, E**. Serotonergic processes passing through the cervical connective (labeled using *Trh⋂tsh*) are active during walking. **(D)** A single animal’s forward velocity with overlaid boxes showing defined walking bouts. **(E)** While tdTomato baseline signal (purple line) is not affected by walking bouts, the calcium signal (green line) in these serotonergic processes rises during walking bouts (gray boxes). **F**. Fluorescent signal in these processes rises with the onset of walking bouts. For each animal, all walking bouts were synchronized around their onset, and an average was taken (between 80 and 130 walking bouts per animal). Plotted is the average of all animals (N=5) with a 95% confidence interval representing the spread between animals.

**Figure 3.**
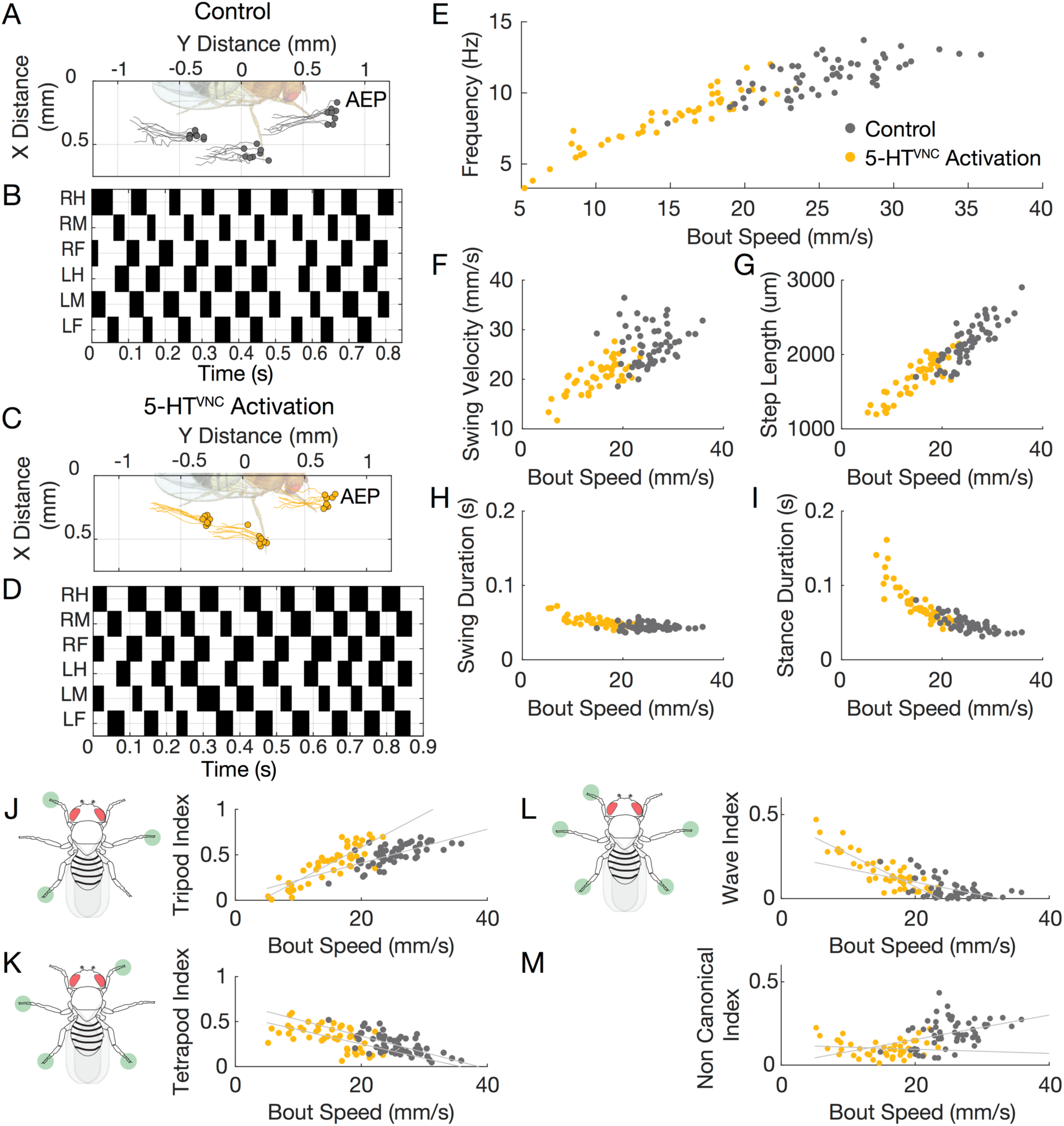
Activation of 5-HT^VNC^ neurons extrapolates most locomotor kinematics. **A-D**. Representative data from speed-matched slow (19 mm/s) walking bouts show that activation of 5-HT^VNC^ neurons does not disrupt locomotor coordination. Footfalls (filled circles) and stance traces (lines) for all steps taken by the left front, middle, and hind legs show foot touchdown placement is consistent over time and stance traces are relatively straight in both control animals (*Trh⋂tsh>csChrimson* grown on food lacking ATR) (A) and animals where 5-HT^VNC^ neurons have been activated (*Trh⋂tsh>csChrimson* fed with ATR) (C). Step trace for each leg during a walking bout for control (B) and experimental (D) animals. Stance phase is indicated in white and swing phase in black. The checkerboard pattern is consistent with a highly-coordinated walking gait. **E-I**. Extrapolation of step parameters upon activation of 5-HT^VNC^ neurons. The relationships between speed and frequency (E), swing velocity (F), step length (G), swing duration (H), and stance duration (I) are shifted in a manner that is consistent with normal walking speeds. N = 47 bouts from 10-23 animals for *Trh⋂tsh>csChrimson* ATR+ (yellow circles). N=56 bouts from 12-30 animals for *Trh⋂tsh>csChrimson* ATR-(gray circles). **J-M**. Activation of 5-HT^VNC^ neurons modifies the relationship between speed and gait selection. Activation of 5-HT^VNC^ neurons increases wave (L) and tetrapod (K) gait utilization while decreasing time spent using tripod (J) gait. There is a low frequency of non-canonical gait conformations upon activation (M). p<.001 for speed by ATR interaction effect in multivariable model for tripod index and wave index. N = 47 bouts from 10-23 animals for *Trh⋂tsh>csChrimson* ATR+ (yellow circles). N=56 bouts from 12-30 animals for *Trh⋂tsh>csChrimson* ATR-(gray circles).

### Flies with activated 5-HT^VNC^ neurons walk in a coordinated manner

Slower walking speeds could be the result of poor coordination or, alternatively, controlled adjustments of gaits, which occur when flies walk upside down or carry an additional load [7]. To distinguish between these two possibilities, we analyzed fly gaits using the Flywalker assay [10]. This system uses frustrated total internal reflection to visualize an animal’s footprints during a walking bout and custom software to analyze these footprints, generating an array of kinematic measurements (Figure S3A) [10].

Using this assay, we find that activation of 5-HT^VNC^ neurons results in highly coordinated walking patterns. Representative traces of an individual’s footprints during a walking bout show that activation of these neurons does not perturb stereotyped foot placement or interfere with the straightness of the stance phase (Figure 3A and C). Step and stance traces show that these animals also use highly coordinated gaits, suggesting that interleg coordination is intact (Figure 3B and D). In fact, compared to control flies, 5-HT^VNC^ neuron activation results in more precise foot placement at the onset and offset of each stance phase, suggesting that the walking behavior of these animals is more constrained compared to control animals (Figure S3B).

In wild type flies, most locomotor parameters are highly correlated with speed, shifting as animals walk faster or slower. Activation of 5-HT^VNC^ neurons induces kinematic shifts that extrapolate these relationships to speeds not normally accessed by wild type flies. For example, as animals walk slower, their step cycle frequency decreases, they take longer steps, and slow the velocity of their swinging legs. These shifts are accompanied by a shift in the step duty cycle, as stance duration increases while swing duration remains largely unchanged [10]. These relationships are maintained and extended into the slower speed range when 5-HT^VNC^ neurons are activated (Figure 3E-I).

In contrast to the kinematic parameters described above, the relationship between gait and speed is not a simple extrapolation upon 5-HT^VNC^ neuron activation. For example, as animals walk more slowly, their preferred gait shifts from the three-legged tripod gait to the more stable tetrapod and wave gaits [8,10]. Upon activation of 5-HT^VNC^ neurons, the slope of a subset of these relationships (tripod and wave gait in particular) shifts (Figure 3J and L). Thus, in addition to maintaining the relationships between speed and most kinematic parameters, serotonin alters the relationship between particular types of gait choice and walking speed.

### Inhibition of 5-HT^VNC^ neurons increases walking speed

Although the experiments described above demonstrate that activation of 5-HT^VNC^ neurons causes flies to walk more slowly, gain-of-function experiments such as these cannot address if and in what situations these neurons are normally used to modulate walking speed. To begin to address this question, we expressed the inward rectifying potassium channel Kir2.1 to constitutively inactivate 5-HT^VNC^ neurons [59]. Although neurons were inactivated throughout development and adulthood, we did not observe a change in the number or anatomy of these neurons in the VNC, suggesting that their development is not significantly affected (Figure S2H and I).

Consistent with the activation phenotype, inhibition of 5-HT^VNC^ neurons causes animals to walk faster (Figure 2B) and increases their angular velocity (Figure S2E and G). Inhibiting these neurons also increases the percentage of time that animals spend walking (Figure S2E).

The shifts in velocity produced by either optogenetic activation or constitutive inhibition of 5-HT^VNC^ neurons mirror each other (Figure 2C). In both cases, ∼12 mm/s is the boundary between velocities that are lost and gained due to 5-HT^VNC^ neuron manipulation (Figure 2C). These complementary shifts in walking speed suggest that serotonin release in the VNC may serve as a switch to regulate behavioral state.

### 5-HT^VNC^ neurons are active in walking flies

The opposing effects on speed when 5-HT^VNC^ neurons are activated or inhibited suggest that the activity of these neurons will co-vary with walk-stop transitions and velocity changes during baseline walking. To test this prediction, we performed functional calcium imaging of a subset of 5-HT^VNC^ axons within the VNC while flies walked on a spherical treadmill (Figure S4A) [60]. To record the largest functional signals, we focused on axons in the neck connective, which are likely derived from the subset of ascending 5-HT^VNC^ neurons that target the brain (Figure S4B-D).

Activity in these fibers is highly correlated with walking (Figure 2D and E). Fluorescence signals from these cells rise dramatically at the onset of each walking bout (Figure 2F). These responses are not nearly as large when animals perform other motor behaviors, like proboscis extension or grooming (Figure S4E). These results suggest that at least a subset of 5-HT^VNC^ neurons are specifically active when flies walk, and are not generically active during all legged motor behaviors. We also find that the activity of these serotonergic processes correlates with the average speed of the walking bout, suggesting that these neurons may become more active when animals walk faster (Figure S4F). These observations are particularly interesting in the context of our other behavioral data, which show that walking speed can be shifted faster or slower by manipulating 5-HT^VNC^ neural activity. Thus, baseline walking speed correlates with but is also sensitive to 5-HT^VNC^ neural activity. Taken together, these observations suggest that one role for serotonin release in the VNC may be to dampen walking speed, a conclusion that we further test below.

### Serotonin slows baseline walking in multiple contexts

Results from optogenetic activation and recordings can be reconciled by a model whereby the VNC serotonergic system is used to regulate walking speed: when the system is activated, flies walk more slowly and when the system is silenced, flies walk faster. To test this model, we asked if 5-HT^VNC^ neural activity is required when flies naturally alter their walking speeds. For example, flies normally walk at different speeds depending on ambient temperature, body orientation, nutritional status, and in response to mechanosensory stimulation [2,7,61,62]. Surprisingly, animals in which 5-HT^VNC^ neurons were silenced were still able to adjust their speed in the same direction as wild type flies in all of these contexts (Figure 4A). Moreover, regardless of the context, animals in which these neurons are silenced walk faster than controls. Thus, animals do not require 5-HT^VNC^ neuron activity to modulate their speed in response to different environmental contexts (e.g. temperature), or internal state (e.g. hunger). Furthermore, these data suggest that this system is used in a context-independent manner to dampen walking speed.

**Figure 4.**
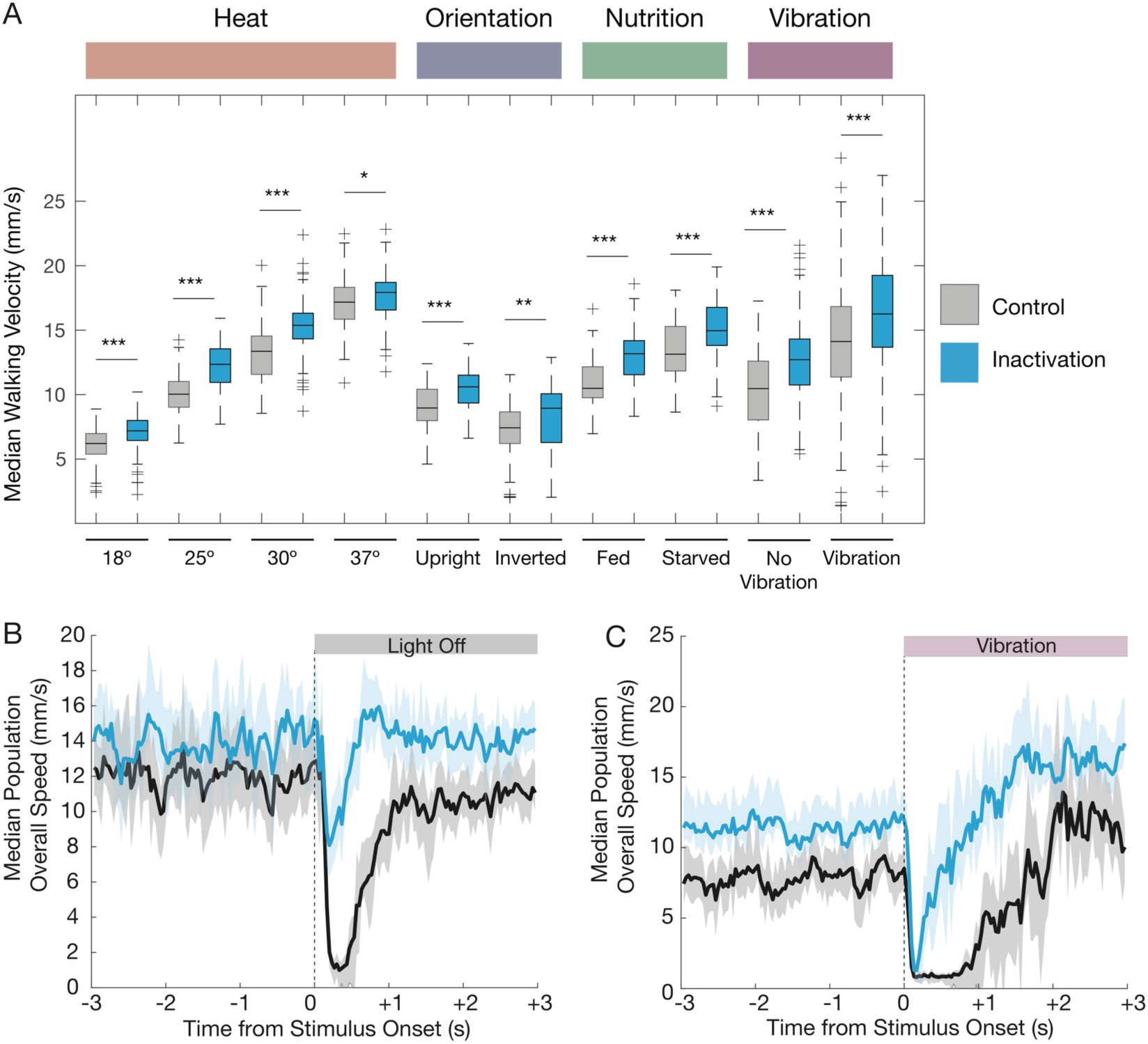
Context-independent and context-specific roles of serotonin in locomotion. **A**. Silencing 5-HT^VNC^ neurons (*Trh⋂tsh>Kir2*.1) causes an increase in walking speed compared to genetically background matched non-Gal4 controls (*w*^*1118*^ *⋂tsh>Kir2*.*1*) across a diversity of behavioral contexts including variations of temperature, orientation, nutritional state, and vibration stimuli. For each condition, genotypes were compared using a Kruskal-Wallis test, ***p<.001, **p<.01, *p<.05. 18 °C– N=130 per genotype. 25 °C – N = 120 per genotype. 30 °C – N = 120 per genotype. 37 °C – N= *w*^*1118*^ (120) *Trh* (119). Upright – N = 90 per genotype. Inverted – N=90 per genotype. Fed – N=86 per genotype. Starved – N = 86 per genotype. Vibration – N= *w*^*1118*^ (167) *Trh* (166). **B-C**. Silencing 5-HT^VNC^ neurons changes the immediate behavioral responses to sudden contextual changes. When lights switch from on to off (B), control animals (*w*^*1118*^ *⋂tsh>Kir2*.*1*, shown in black) show a brief behavioral pause (indicated by arrow) and then resume activity. However, when 5-HT^VNC^ neurons are silenced, animals slow their speed but do not fully pause. In response to the onset of vibration (C) control animals stop, pause and then accelerate speed. When 5-HT^VNC^ neurons are silenced (*Trh⋂tsh>Kir2*.1, shown in blue), animals pause but re-accelerate more quickly than controls. Shaded areas show 95% confidence intervals. For light experiments, N= *w*^*1118*^ (150) *Trh* (140); for vibration experiments N= *w*^*1118*^ (167) *Trh* (166).

### VNC serotonin release is required on a fast time scale to respond to rapid contextual changes

Another scenario in which animals may benefit from slowing down is when they are startled. In mammals, stereotyped startle behaviors occur in response to a wide variety of sensory stimuli – acoustic, tactile, and vestibular. These responses take place on sub-second time scales, involve simultaneous contraction of muscles throughout the body, and are similar irrespective of the initiating stimulus [63-65]. Like mammals, *Drosophila* display stereotyped responses to threatening looming stimuli, beginning with an initial freezing period lasting less than a second before escape behaviors are initiated [66,67]. Because these startle responses are contextually independent and have been shown to be mediated in part by serotonin in mammals [68], we next asked whether 5-HT^VNC^ neurons are required for these responses in *Drosophila*.

We tested this prediction using two different startle-inducing paradigms: (i) one in which flies abruptly experience total darkness (‘blackout paradigm’) and (ii) one in which flies suddenly experience strong mechanical stimulation, such as an intense vibration (‘earthquake paradigm’) [62]. In both scenarios, control animals show a two-tiered response to these abrupt changes (Figure 4B and C): First, animals rapidly come to a nearly complete stop. Then, they pause before resuming a behavior that is appropriate for the new context. For both the blackout and earthquake paradigms, control animals stop within the first 0.25 seconds, pause for about a second, and only then resume walking behavior (Figure 4B and C, Figure S5B and D). Animals lacking the ability to release serotonin in the VNC are deficient in these initial responses, but are still able to eventually achieve context-appropriate walking speeds (Figure 4A). Moreover, consistent with our earlier analyses, flies unable to activate 5-HT^VNC^ neurons walk faster compared to control flies, both before and after the change in their environment (Figure 4B and C).

Taken together, these results demonstrate that the VNC serotonergic system is not required for animals to modify their walking speed in response to changes in the environment or shifts in internal state. In addition, they suggest that this system is needed for an immediate and stimulus-independent response when flies are startled.

### Different serotonin receptor mutants alter the startle response in different ways

All five serotonergic receptors in *Drosophila* – 5-HT1A, 5-HT1B, 5-HT2A, 5-HT2B, and 5-HT7 – are G-protein coupled receptors (GPCRs) [69-72]. Like their mammalian orthologs, members of each serotonin receptor family (1, 2, and 7) have distinct cellular effects upon activation. Receptors in the 1 family, 5-HT1A and 5-HT1B, act through the G_i_ pathway to inhibit the generation of cAMP, whereas 5-HT7, the only member of the 7 family in *Drosophila*, stimulates the production of cAMP [70,73]. Receptors of the 2 family, 5-HT2A and 5-HT2B in *Drosophila*, act through a the PLC-IP_3_ signaling pathway to increase intracellular calcium [72,74,75]. Together, this diversity of receptors is thought to allow serotonin to produce complex physiological responses that depend on both synaptic connectivity and receptor expression patterns.

Before characterizing the phenotypes of these receptor mutants, we analyzed a mutant of the *Trh* gene (*Trh*^*01*^), which is globally unable to produce serotonin [48]. Reassuringly, *Trh*^*01*^ animals show a similar phenotype to animals in which 5-HT^VNC^ neurons were silenced: the flies walk significantly faster and more frequently than controls, and exhibit a similar startle response in the earthquake paradigm (Figure 5A and B, Figure S5A,C-F). In addition, *Trh*^*01*^ mutant animals walk closer to the edge of the arena compared to control animals, consistent with previous observations [46] (Figure S5A).

**Figure 5.**
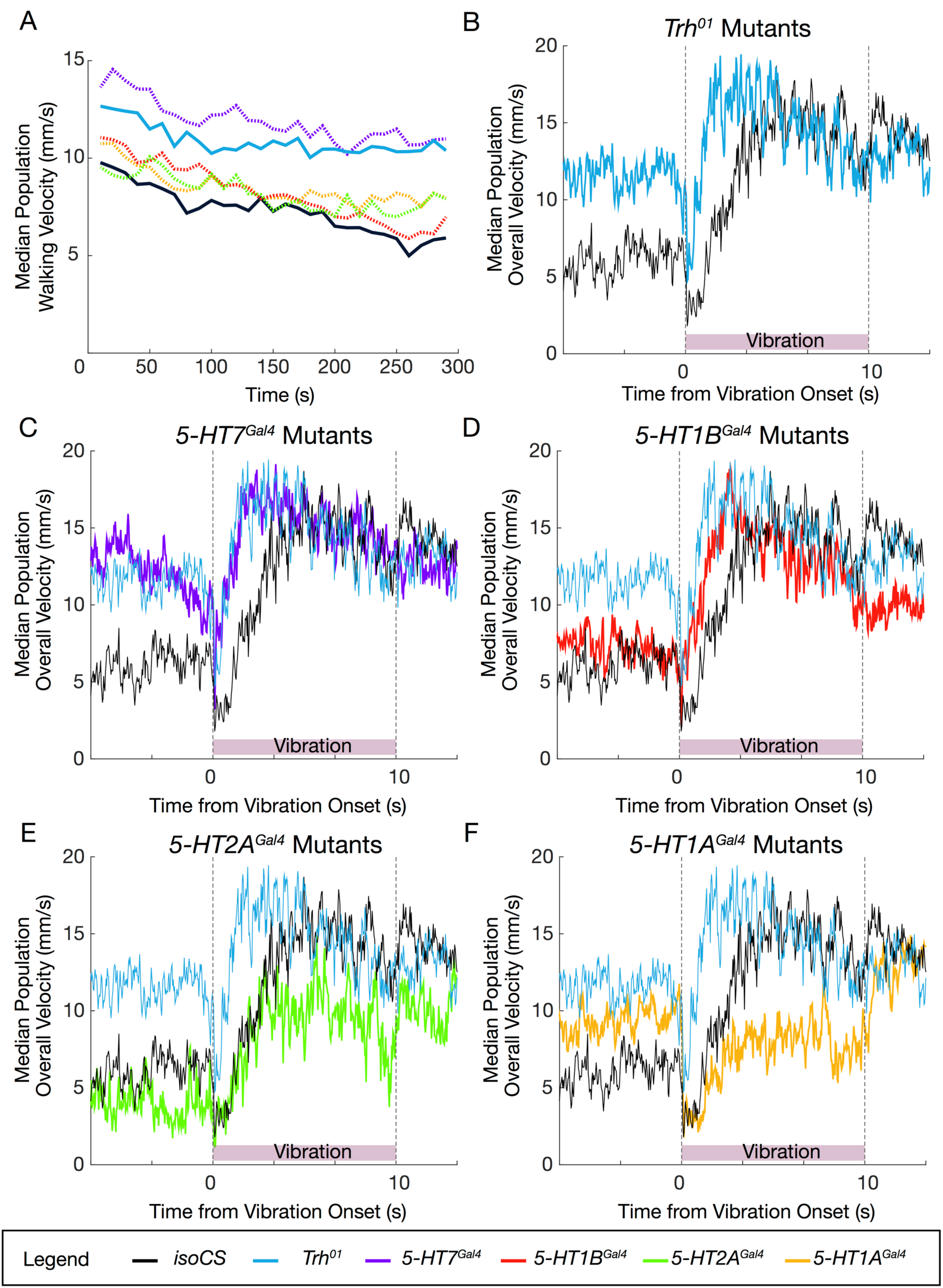
Mutations in *Trh* and select serotonin receptors replicate 5-HT^VNC^ inactivation phenotype. **A**. Median population walking speed binned at ten second intervals for *Trh*^*01*^ mutants (blue), which walk faster than background matched *isoCS* controls (black), consistent with the inactivation experiments. *5-HT7*^*Gal4*^ mutants (purple) replicates this phenotype, but other mutants do not show a baseline increase in walking speed. N=130 *isoCS*, N=130 *5-HT1A*^*Gal4*^, N=120 *5-HT1B*^*Gal4*^, N=100 *5-HT2A*^*Gal4*^, N=120 *5-HT7*^*Gal4*^, N=120 *Trh*^*01*^. **B-F**. Median population walking speed sampled at 30 Hz in response to vibration stimulus. *Trh*^*01*^ mutants (B, blue line) show a blunted and shortened pause in response to novel stimulus. *5-HT7*^*Gal4*^ mutants (C, purple line) and *5-HT1B*^*Gal4*^ mutants (D, red line) show a similar phenotype to *Trh*^*01*^ mutants (blue line). *5-HT2A*^*Gal4*^ mutants (E, green line) and *5-HT1A*^*Gal4*^ mutants (F, yellow line) have a pause phase comparable to controls, but do not accelerate as much in response to the vibration stimulus.

Null receptor mutants *5-HT1A*^*Gal4*^, *1B*^*Gal4*^, *2A*^*Gal4*^, and *7*^*Gal4*^ all increase the percentage of time animals spend walking (Figure S5A) [48]. However, an increase in walking speed is only observed in *5-HT7*^*Gal4*^ mutants (Figure 5A and C), suggesting that 5-HT7 is the primary receptor responsible for mediating the effects of serotonin on walking speed.

We next tested the receptor mutants in the earthquake paradigm. Interestingly, *5-HT7*^*Gal4*^ and *5-HT1B*^*Gal4*^ mutants closely phenocopy the startle response seen in *Trh*^*01*^ mutants and also in the 5-HT^VNC^ inactivation experiments (Figure 5C and D, Figure S5E-H). By contrast, although *5-HT1A*^*Gal4*^ and *5-HT2A*^*Gal4*^ mutants speed up at the same rate as controls, they exhibit a sustained decrease in their final target speed in response to this stimulus (Figure 5E and F, Figure S5E, I, and J).

These data are consistent with the idea that different receptors influence distinct aspects of the startle response. Notably, mutation of receptors that are predicted to have opposing effects on cAMP production, such as 5-HT1A and 5-HT7, can result in similar phenotypes. Further, some receptor mutants exhibit phenotypes that are not seen in *Trh*^*01*^ mutant animals. These complex changes in locomotor behavior may be explained by the differential expression of serotonin receptors in key components of the locomotor circuit.

### Serotonin receptors are expressed in distinct cell types

The different effects on walking observed in flies mutant for different serotonin receptors suggest that, in addition to distinct biochemical properties, they also have different expression patterns within the locomotor circuit. To identify neurons that express these receptors, we used gene and protein trap Gal4 lines from the MiMIC library to drive expression of a GFP reporter in the pattern of each receptor subtype [76] (Figure S6G). Each receptor line drove expression in many neurons both within the brain and the VNC (Figure 6A-E). Many of these are uncharacterized interneurons that cannot yet be functionally studied. However, each serotonin receptor is also expressed in distinct subsets of leg motor and sensory neurons. In particular, while members of the 5-HT1 family are predominantly expressed in mechanosensory neurons throughout the leg, 5-HT2 and 5-HT7 receptors are expressed in proximal-targeting flexor and extensor motor neurons (Figure S6A-F), proprioceptive, and distal sensory neuron populations (Figure 6F-K). Thus, serotonin release in the VNC is likely to differentially affect these components of the locomotor circuit, ultimately contributing to the observed changes in behavior.

**Figure 6.**
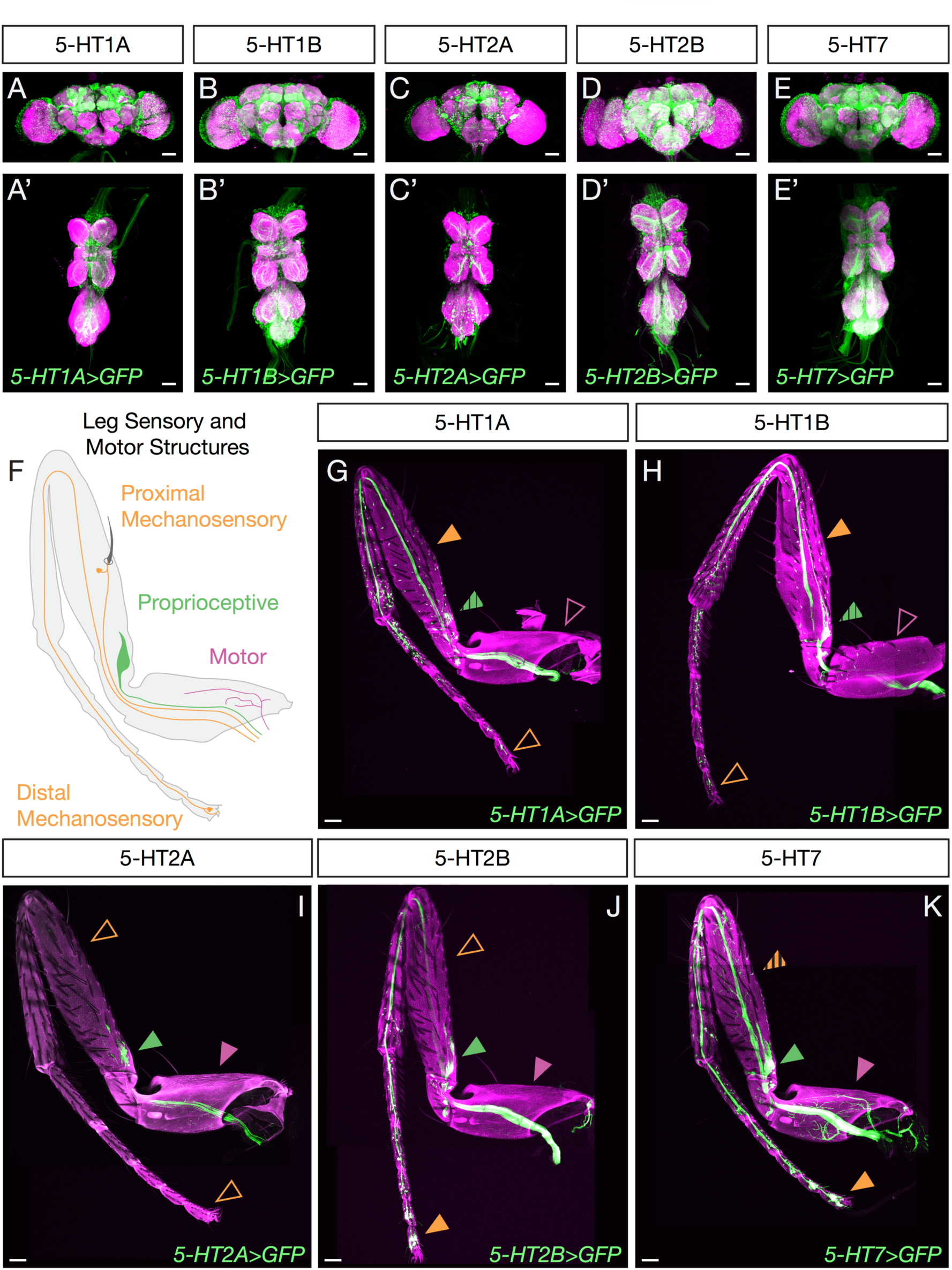
Serotonin receptors are differentially expressed in locomotor circuitry components, including sensory and motor neurons. **A-E**. Maximum intensity projections show Gal4-driven expression of serotonin receptors in both the brain and VNC. All scale bars are 50 µm. **F**. Schematic of sensory and motor neuron populations innervating the adult leg. **G-K**. Maximum intensity projections show Gal4-driven expression of serotonin receptors in neuronal processes in the adult T1 leg. Each receptor is expressed in a distinct pattern in sensory-motor components. While some are expressed in motor neurons (purple arrows, hatched arrows indicate limited or weak expression) and proprioceptive neurons (green arrows), others are not. All receptors are expressed in a subset of mechanosensory neurons (orange arrows), but some are preferentially expressed in proximal or distal leg segments. All scale bars are 50 µm.

## Discussion

Walking is highly stereotyped, consisting of a small number of well-defined gaits, each with its own set of characteristic kinematic parameters. However, these highly stereotyped gaits must also be flexible, to adapt to a wide variety of environments, complex terrains, and novel situations. How nervous systems manage to orchestrate behaviors that are simultaneously stereotyped and flexible is not well understood. Here we show in the fly that (i) serotonergic VNC neural activity modulates walking speed in a context-independent manner and (ii) these neurons play a critical role in a fly’s ability to modulate its walking behavior in response to the sudden onset of a startling stimulus. Moreover, the multiple roles the serotonergic system plays are mediated by distinct receptors, which have different biochemical properties and are expressed in unique subsets of locomotor circuit components.

### A common role for serotonin in modulating walking speed across species

A central finding of our study is that serotonergic neuron activity in the VNC modulates baseline walking speed. This finding strongly parallels previous observations in the motor systems of other organisms. For example, activity of serotonergic neurons in the cat brainstem is correlated with motor behavior and walking speed [12,77,78]. Additionally, in vertebrates as diverse as the lamprey and mouse, serotonin induces an increase in step cycle period, slowing locomotor rhythms (reviewed in [42]). One recent study in mice showed that activation of the dorsal raphe nucleus – a key serotonergic brain region – produces rapid suppression of spontaneous locomotion and locomotor speed, while showing minimal effect on kinematic parameters, such as gait, or on non-locomotor behaviors such as grooming [79]. These parallels to our results suggest that the modulatory role of the serotonergic system in regulating locomotor speed is remarkably conserved across the animal kingdom.

### Serotonergic modulation of the startle response

In our experiments, we consistently observed that the onset of a startling stimulus (be it visual or mechanosensory), induces a brief period of pausing behavior in wild type flies. We hypothesize that these behavioral pauses are similar to startle responses seen in both mammalian systems and insects, which also trigger a pause phase before animals embark on an appropriate behavioral action [63,64,67,68,80-82]. While the importance of these pausing behaviors remains poorly understood, it may be that they allow animals to collect additional sensory information before they select an appropriate response to the startling stimulus.

We find that inactivation of the VNC serotonergic system does not completely abolish this startle response, suggesting that this system is not the primary driver of this behavior, but instead serves to modulate the latency of the response following an initial freeze phase when animals stop. A role for serotonin in the startle response appears to be conserved: In mammals, the absence of serotonin, due to the lesion of key serotonergic brain regions or pharmacological blockade, is generally associated with an increase in the intensity of startle responses [83,84]. Although this may seem counter to our results, other studies have shown that serotonin increases startle responses when injected directly into the lumbar spinal cord [85-88]. Thus, serotonin may play distinct roles in the forebrain and in the spinal cord. Together with our results, we suggest that the role of spinal cord/VNC serotonin release is to extend the latency period and/or amplify the startle response.

### Serotonin acts differentially on sensory and motor circuitry to modulate walking

Based on the distribution of receptors in locomotor circuitry components, we can formulate a preliminary model of how serotonergic action may be working to modulate *Drosophila* locomotor circuits both at the level of sensory input and motor output. We note, however, that this model is likely incomplete as we cannot as of now incorporate the role that local interneurons that also express 5-HT receptors play in modulating locomotion behavior. Nevertheless, because the primary receptors expressed in motor neurons are 5-HT7 and 5-HT2B, which have been shown to upregulate the production of cAMP and facilitate calcium entry [72,89], we hypothesize that the phenotypes we observe are due at least in part to serotonin action on these cells.

There is ample evidence in the literature to support this role for serotonin both in rodent models as well as in human studies (reviewed in [90-92]). While increased motor output is usually correlated with increased, not decreased, speed [93], increased muscle output has also been shown to be required in humans to navigate complex terrain, and may be playing a similar role in the fly [94,95]. In addition, it is noteworthy that the motor neurons expressing these receptors target both flexor and extensor muscles in the coxa and femur, two proximal leg segments (Figure S6A-F). These observations suggest that serotonin acting on these motor neurons may result in co-contraction, a mechanism that facilitates joint stability in the face of a complex environment and also during the preparatory phase for certain escape behaviors [66,94-99]

In addition to a potential role in motor neurons, serotonin receptors are expressed in distinct classes of leg sensory neurons that target the leg neuropils of the VNC. We hypothesize that this distribution of receptors serves to shift the balance of sensory information in response to serotonergic input. Based on the known downstream signaling properties of these receptors, we predict that increased levels of serotonin in the VNC would amplify proprioceptive and distal sensory inputs at the expense of more proximal sensory information. These shifts in sensory processing may also contribute to increased stability and might be useful in other contexts where slow walking is preferred, such as navigating complex terrains where improved sensory information might be beneficial.

Considering the broad expression of serotonergic receptors in sensory organs, it is interesting that one of the behavioral roles of serotonin we identified is its ability to mediate the response to vibration. Vibration is sensed by the chordotonal organ, and our expression analysis reveals that serotonin receptors are expressed to different extents in chordotonal neurons [100-102]. Together, these observations suggest that modulation of sensory information as it is entering the VNC plays a key role in how serotonin modulates the response to a vibration stimulus. In addition, the observation that serotonin release in the VNC affects the response to both the earthquake and blackout paradigms similarly, which are perceived by two very different sensory systems, suggests that this neuromodulator is controlling downstream locomotor components, such as motor neurons, that are shared by both systems.

### Experimental Procedures

#### Fly husbandry

Unless otherwise described, flies were maintained at 25° C on dextrose cornmeal food using standard laboratory techniques. Crosses used for behavioral experiments were flipped every 2-3 days to prevent overcrowding. For all arena experiments, flies were maintained on Nutrifly German Sick food (Genessee Scientific 66-115) in an incubator humidified at 60% with a 12h:12h light:dark cycle. As animals eclosed, females of the appropriate genotype were collected under CO_2_ anesthesia every 2-3 days. For non-optogenetic experiments, flies were collected onto Nutrifly Food without any additive. For optogenetic experiments, flies were collected onto Nutrifly food supplemented with either .4 mM ATR or an equal concentration of solvent alone (DMSO for Flywalker experiments, 95% EtOH for arena experiments). Animals were aged in the dark (for optogenetic experiments) or on the same light:dark cycle for 2-3 more days at 25° C before being assayed.

#### Fly Strains

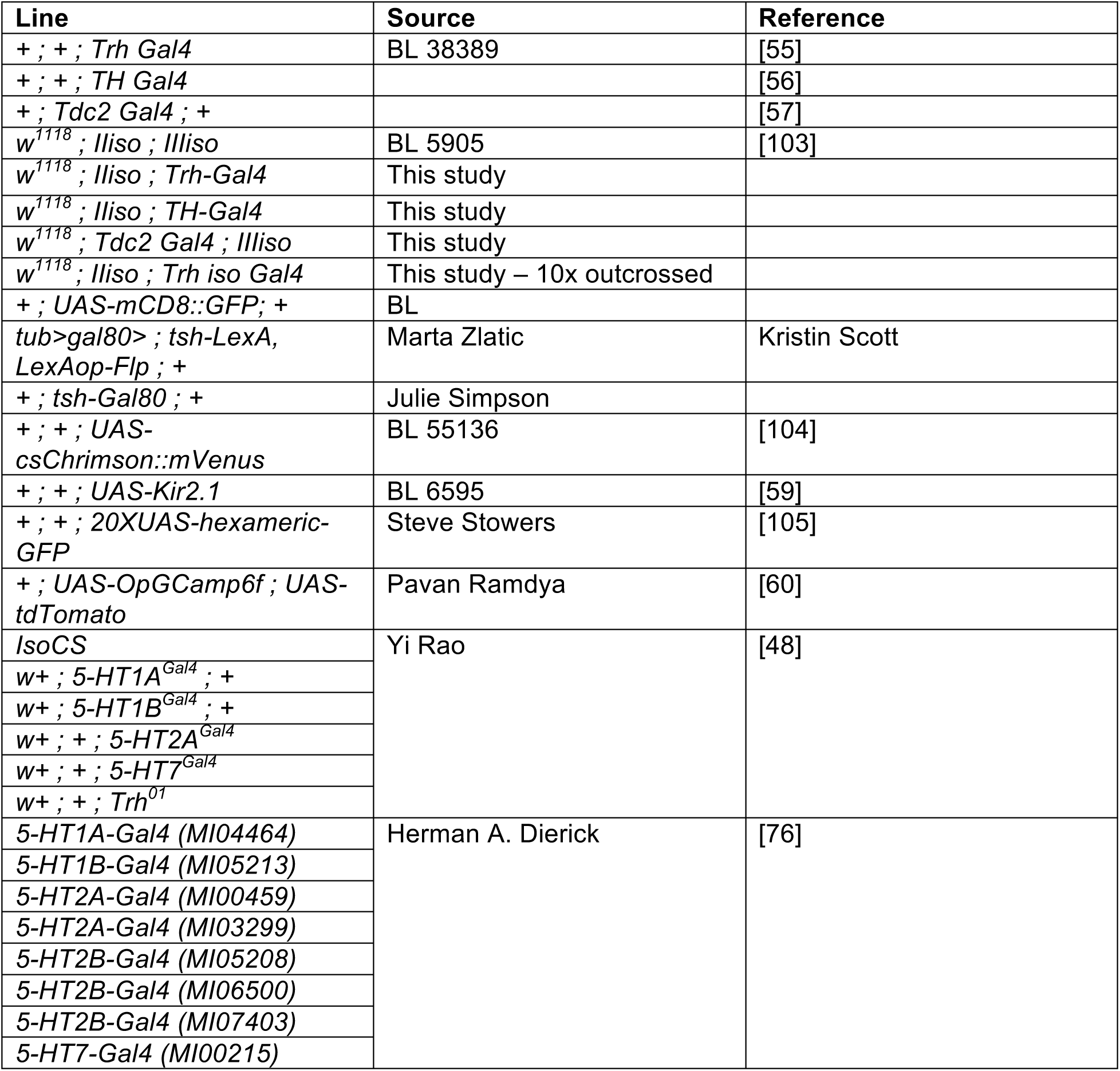
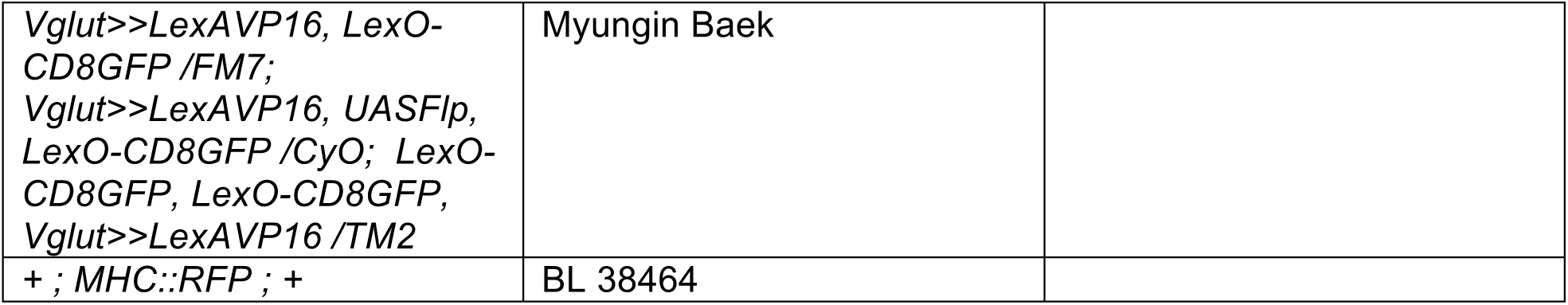

#### Genotypes with associated Figures

| Experimental Line | Main Figure | Supplementary |
| --- | --- | --- |
| *+ /+ ; tsh Gal80 / UAS mCD8 GFP ; Trh Gal4 / +* | 1B, B', B'' |  |
| *tub>gal80> / + ; tsh LexA, lexAop Flp / + ; Trh Gal4 / UAS mCD8 GFP* | 1C, C', C'' |  |
| *+ /+ ; tsh Gal80 / +; UAS mCD8 GFP, TH Gal4 / +* | 1D, D', D'' |  |
| *tub>gal80> / + ; tsh LexA, lexAop Flp / + ; TH Gal4, UAS mCD8 GFP / +* | 1E, E', E'' |  |
| *+ /+ ; tsh Gal80 / Tdc2 Gal4 ; UAS mCD8 GFP / +* | 1F, F', F'' |  |
| *tub>gal80> / + ; tsh LexA, lexAop Flp / Tdc2 Gal4 ; UAS mCD8 GFP / +* | 1G, G', G'' |  |
| *tub>gal80> / w<sup>1118</sup> ; tsh LexA, lexAop Flp / isoII ; UAS csChrimson::mVenus / isoIII* | 1H<br>2A, C | S2D, F |
| *tub>gal80> / w<sup>1118</sup> ; tsh LexA, lexAop Flp / isoII ; UAS csChrimson::mVenus / Trh Gal4* | 1H<br>3A-M | S3 B<br>S4 D |
| *tub>gal80> / w<sup>1118</sup> ; tsh LexA, lexAop Flp / isoII ; UAS csChrimson::mVenus / TH Gal4* | 1H |  |
| *tub>gal80> / w<sup>1118</sup> ; tsh LexA, lexAop Flp / Tdc2 Gal4 ; UAS csChrimson::mVenus / isoIII* | 1H |  |
| *tub>gal80> / w<sup>1118</sup> ; tsh LexA, lexAop Flp / isoII ; UAS csChrimson::mVenus / Trh iso Gal4* | 2A, C | S1<br>S2D, F, H, I |
| *tub>gal80> / w<sup>1118</sup> ; tsh LexA, lexAop Flp / isoII ; UAS Kir2.1::GFP / isoIII* | 2B and C<br>4A-C | S2E and G |
| *tub>gal80> / w<sup>1118</sup> ; tsh LexA, lexAop Flp / isoII ; UAS Kir2.1::GFP / Trh iso Gal4* | 2B and C<br>4A-C | S2E, G, H, I<br>S5D and F |
| *tub>gal80>+/+ ; tsh-LexA, lexAop-FLP/UAS-opGCaMP6f; Trh-Gal4/UAS-tdTomato* | 2D-F | S4B, C, E, F |
| *isoCS* | 5A-F | S5A,C, E |
| *W+ ; 5-HT1A<sup>Gal4</sup> ; +* | 5A, F | S5A,C,J |
| *W+ ; 5-HT1B<sup>Gal4</sup> ; +* | 5A, D | S5A,C,H |
| *W+ ; + ; 5-HT2A<sup>Gal4</sup>* | 5A, E | S5A,C,I |
| *W+ ; + ; 5-HT7<sup>Gal4</sup>* | 5A, C | S5A,C,G |
| *W+ ; + ; Trh<sup>01</sup>* | 5A-F | S5A,C-J |
| *5-HT1A-Gal4 (MI04464) / + ; 20X-UAS hexameric GFP / +* | 6A,G | S6G |
| *5-HT1B-Gal4 (MI05213) / + ; 20X-UAS hexameric GFP / +* | 6B,H | S6G |
| *+/+ ; 5-HT2A-Gal4 (MI00459) / 20X-UAS hexameric GFP* | 6C,I | S6G |
| *+/+ ; 5-HT2B-Gal4 (MI05208) / 20X-UAS hexameric GFP* | 6D, J | S6G |
| *+/+ ; 5-HT7-Gal4 (MI0215) / 20X-UAS hexameric GFP* | 6E, K | S6G |
| *+/+ ; 5-HT2A-Gal4 (MI03299) / 20X-UAS hexameric GFP* |  | S6G |
| *+/+ ; 5-HT2B-Gal4 (MI06500) / 20X-UAS hexameric GFP* |  | S6G |
| *+/+ ; 5-HT2B-Gal4 (MI07403) / 20X-UAS hexameric GFP* |  | S6G |
| *Vglut>>LexA, LexO-CD8GFP / +; Vglut>>LexA, UASFlp, LexO-CD8GFP / +; LexO-CD8GFP, LexO-CD8GFP, Vglut>>LexAVP16 / 5-HT2B-Gal4 (MI05208)* |  | S6A |
| *Vglut>>LexA, LexO-CD8GFP / +; Vglut>>LexA, UASFlp, LexO-CD8GFP / +; LexO-CD8GFP, LexO-CD8GFP, Vglut>>LexAVP16 / 5-HT7-Gal4 (MI0215)* |  | S6B |
| *Vglut>>LexA, LexO-CD8GFP / +; Vglut>>LexA, UASFlp, LexO-CD8GFP / MHC-RFP; LexO-CD8GFP, LexO-CD8GFP, Vglut>>LexAVP16 / 5-HT2B-Gal4 (MI05208)* |  | S6C |
| *Vglut>>LexA, LexO-CD8GFP / +; Vglut>>LexA, UASFlp, LexO-CD8GFP / MHC-RFP; LexO-CD8GFP, LexO-CD8GFP, Vglut>>LexAVP16 / 5-HT7-Gal4 (MI0215)* |  | S6D |

#### Immunostaining brain and VNC

Brains and VNCs were dissected in phosphate buffered saline with 0.3% Triton (PBST) and fixed in 4% Paraformaldehyde (PFA) for 20 minutes. Samples were washed five times for 20 minutes in PBST with 0.1% Bovine serum albumin (BSA), and then blocked in PBST-BSA for one hour at room temperature, or overnight at 4° C. Samples were incubated with primary antibody diluted in PBST-BSA overnight at 4° C, and washed five times 20 minutes with PBST-BSA the next day. Samples were then incubated in secondary antibody diluted in PBST-BSA overnight at 4° C. The next day, samples were washed five times for 20 minutes in PBST, and then the liquid was replaced with Vectashield and samples were incubated overnight prior to mounting. Brains and VNCs from the same animals were mounted together, with the ventral surface of the VNC and the anterior surface of the brain facing up.

Validation of expression patterns during two photon experiments were performed as described in [60].

#### Antibodies

Antibodies:
| Antibody | Source | Concentration | Animal | Figure | SFig |
| --- | --- | --- | --- | --- | --- |
| Vglut | Aaron DiAntonio | 1:10,000 | rb |  | S1J |
| 5-HT | Sigma | 1:1000 | rb |  | S1E, S4D' |
| TH | Immunostar | 1:1000 | m |  | S1F |
| ChAT | DHSB | 1:500 | m |  | S1I |
| Brp-c | DHSB | 1:20 – 1:100 | m | 1B-G, 6A-E | S4B |
| GABA | Sigma | 1:1000 | rb |  | S1H |
| Tdc2 | Cova Labs | 1:200 | rb |  | S1G |
| dsRed | Takara Bio | 1:1000 | rb |  | S4 B |

| Animal | Fluorophore | Concentration | Source |
| --- | --- | --- | --- |
| r | Alexa 488 | 1:500 | Invitrogen |
| rb | Alexa 555 | 1:500 | Invitrogen |
| m | Alexa 647 | 1:500 | Invitrogen |
| Goat anti rb | Cy3 | 1:400 | Jackson |
| Goat anti m | Alexa 633 | 1:400 | Thermofisher |

#### Confocal Imaging

Mounted brains and VNCs were imaged on a Leica TCS SP5 confocal at 20X magnification with a resolution of 1024 x 512 pixels, and at a scanning rate of 200 Hz and 3x averaging. Sections were taken at 1 um increments. Laser power and detector gain were maintained constant for the brain and VNC of the same animals, but were adjusted for optimal signal between animals.

Imaging of fixed samples following two photon live imaging experiments was performed on a Zeiss LSM 700 Laser Scanning Confocal Microscope at 20X magnification and 2X averaging, with a 0.52 × 0.52 um pixel size. Z sections were taken at 1 um intervals. As described in [60].

#### Cell Counting and Quantification

Images were analyzed in Fiji [106]. For quantification of the number of cells driven by *Trh-Gal4* in the brain and VNC, mVenus positive and 5-HT positive cell bodies were counted from five or more individual animals.

#### Leg dissection, imaging, and image processing

To prepare legs for imaging, fly heads and abdomens were removed, and thoraces with legs attached were fixed overnight in 4% PFA at 4° C. Carcasses were washed 5 times with 0.03% PBST, and then place in Vectashield overnight before legs were mounted. Imaging was performed on a Leica SP5 confocal at 20X and 1024 x 1024 pixel resolution with 3x averaging, with sections taken at steps of 1 um. Two PMT detectors were set to capture green fluorescent signal and the green autofluorescence of the cuticle. Laser power was adjusted independently for each line to achieve optimal visualization of structures. Images were processed in Fiji [106]. Autofluorescence was subtracted from the green channel to allow for clearer visualization of leg structures.

## Behavioral Systems

### Arena Experiments

#### Hardware

The skeleton of the system was built of 80-20 bars and acrylic plates and the arena itself was machined out of polycarbonate to the specifications published in [107]. The polycarbonate plastic arena was embedded in an aluminum plate to maintain a level surface. During experiments the arena was covered with an acrylic disc with a small hole for mouth pipetting in flies. The inside of the lid was coated in a thin layer of Fluon (Amazon, B00UJLH12A) to prevent flies from walking on the ceiling.

A Point Grey Blackfly Mono USB3 camera fitted with a Tamron 1/2” F/1.2 IR C-mount lens (B&H photo) was mounted above the arena and connected by USB 3 cable to a System 76 Leopard WS computer running Ubuntu 14.04 LTS. A Kodax 3×3” 89B Opaque IR filter (B&H photo) was placed in front of the camera detector to allow for detection of IR but not visible light.

Backlighting and optogenetic stimulation was provided by a plate of LEDs sitting under the arena. An acrylic diffuser was placed between the lighting plate and the arena. Each plate was designed with two sets of LEDs – one for IR backlighting (ledlightsworld.com SMD3528-300) and one for optogenetic or white light stimulation (superbrightLEDS.com NFLS-x-LC2 in Red or Natural White). These plates were swapped out when experiments required different color LEDs. To allow for detection of the on state of optogenetic lights, an additional IR light was wired in series with each visible light array, and placed within the field of view of the camera.

Each set of lights was powered separately by an Arduino Uno driver, allowing for modulation of light intensity via Pulse Width Modulation (PWM). Commands to set LED brightness and start and end experiments were sent to this driver using a PuTTY terminal and USB serial interface. For all the experiments described here, both IR and visible spectrum LEDs were set at 100% brightness. At the center of the arena this corresponded roughly to intensities of:

| Light On | Percent | Intensity | Wavelength Measured |
| --- | --- | --- | --- |
| Infrared | 100 | .13 mW | 1050 nm |
|  |  | .08 mW | 635 nm |
|  |  | .08 mW | 535 nm |
| Infrared + Red | 100 | .6 mW | 635 nm |
| Infrared + White | 100 | .62 mW | 635 nm |
|  |  | .68 mW | 535 nm |

## Data Acquisition

All behavioral recordings were done during the three-hour morning activity peak. Prior to the experiment, the arena was leveled, the lid cleaned, and a new layer of Fluon applied. For each experiment, videos were recorded of cohorts of ten flies. For each recording session, flies were mouthpipetted into the arena through a small hole and then the arena lid was slid to move the hole out of the field of view. A blackout curtain cover (Thor Labs, BK5) was used to surround the arena, protecting it from any contaminating light.

Experimental protocols were programmed into the Arduino through serial communication via a PuTTY terminal. Videos were recorded at a rate of 30 frames per second and stored in a compressed “fly movie format” using custom software written by Andrew Straw at the University of Freiberg based on work previously described [108].

### Orientationexperiments

For inverted experiments, animals were introduced into the arena set-up when it was upright, and the lid of the arena was taped in place. The entire arena was manually inverted and propped up on two overturned ice buckets. Flies were either recorded upright and then inverted, for six minutes each, or in the opposite order. As no indicator was present to identify the moment of inversion, the first five minutes and the final five minutes of the video were selected as the before and after orientation switch periods.

### Starvation

24 hours prior to behavioral assay, half of the flies were transferred to an empty tube with a wet Kim Wipe. Behavioral recordings were collected as described above and lasted for five minutes.

### Heat

Heated experiments were carried out inside a walk-in temperature-controlled incubator, which was set at either 18, 25, 30, or 37 C and 40% humidity. Flies were introduced to the arena immediately after entering the temperature-controlled room, recording began immediately thereafter and lasted for five minutes.

### Light

For experiments examining responses to light stimuli, flies were first exposed to five minutes of white light, and then a one minute period of darkness.

### Vibration

To provide a vibration stimulus, four 3V haptic motors (1670-1023-ND, Digikey) were attached to the aluminum plate in which the arena sat using 3D printed holders. The motors were wired in series and driven by the same Arduino system driving the arena’s LED lighting array. For all experiments described, vibration was set at 10% power. The protocol for vibration experiments consisted of a brief habituation period (five minutes for inactivation experiments, 30 seconds for mutant experiments) followed by a 10 second vibration pulse and a 110 sec recovery period.

## Tracking

Videos were tracked using the FlyTracker software from the Caltech vision lab [109]. Prior to tracking, pixel to mm conversion was calibrated using an inbuilt GUI. One calibration file was generated for all videos taken on the same day. Background model and thresholds were adjusted to provide optimal recognition of animals and were not standardized between recording sessions. If present, the state of an indicator light was annotated by custom-written MATLAB software.

## Behavioral classifiers

### Jump

Jumps were classified as frames where the velocity of the animal exceeded 50 mm/s.

### Walk

Walking frames were defined using a dual threshold Schmitt trigger filter. Speed thresholds were set at 1 and 2.5 mm/s, and time thresholds were 0.1 s. Walking frames were also specified to be those in which the fly was not already engaged in a jump.

### Stop

Stop frames were classified as any frames where animals were not performing walking or jumping behaviors.

## Parameters

| Tripod | Tetrapod | Wave | Non-Canonical |
| --- | --- | --- | --- |
| 100110 | 011011 | 011111 | All Other |
| 011001 | 011110 | 101111 |  |
|  | 100111 | 110111 |  |
|  | 110110 | 111011 |  |
|  | 101101 | 111101 |  |
|  | 111001 | 111110 |  |

### WalkFrequency

the percent of frames classified as walking during the recording period.

### OverallVelocity

the median of all velocities over the recording period.

### WalkingVelocity

the median of velocities during all frames when the animal is classified as walking.

### MaximumWalkingVelocity

the maximum velocity an animal reaches during walking.

### Angularvelocity

the median value of angular velocity. This parameter takes into account directionality of turning.

### AbsoluteAngularVelocity

the median of the absolute value of angular velocities. This parameter does not take into account directionality of turning.

### DistancefromWall

the median distance from the closest point on the arena wall during the recording period.

### Walkingboutnumber

bouts were defined as contiguous frames of walking (longer than .1 s as specified in the walking classifier). The number of bouts was calculated for the entire recording period.

### Walkingboutduration

the length of each bout was calculated, and the median of all bout lengths was taken for each animal.

### Stopboutnumberandduration

calculated as for walking bouts.

### JumpFrequency

the percent of time that an animal spends in the jump state as defined above.

## Statistics

For optogenetic arena experiments, behavior during the five-minutes before optogenetic lights were turned on was compared to behavior for five minutes of the light on period for each individual, with a 30 second buffer period after the light was turned on to avoid contamination from behavioral reaction to the light itself. For each activation experiment, we recorded behavior from both experimental (*Trh⋂tsh>csChrimson*) and control (*w*^*1118*^ *⋂tsh>csChrimson*) flies. Prior to these experiments, the *Trh Gal4* line was outcrossed 10 times to our isogenized *w*^*1118*^ control to ensure experimental and control backgrounds were genetically matched. For each genotype, we analyzed data from flies that had been fed with ATR, the required co-factor for optogenetic activation, and flies from the same cross that had been fed on food depleted of ATR. Figures show comparisons between ATR+ control and experimental animal behavior, as we found these populations had the most similar light off behavior pattern. However, significant differences in parameters are consistent even when all controls are included in the analysis.

For constitutive inhibition arena experiments, behavior during the five-minute light off period was analyzed for experimental (*Trh⋂tsh>Kir2*.*1*) and control (*w*^*1118*^ *⋂tsh>Kir2*.*1*) flies fed on the same ATR negative food we used for optogenetic experiments. Prior to these experiments, the *Trh Gal4* line was outcrossed 10 times to our isogenized *w*^*1118*^ control to ensure experimental and control backgrounds were genetically matched.

All analysis on data from arena experiments was performed in MATLAB using custom-written scripts. For normally distributed data, groups were compared by t-test. For non-normal data, groups were compared using Kruskal-Wallis analysis with Dunn-Sidak multiple correction testing when multiple groups were being compared. To compare changes in velocity distribution, bootstrapping was used to estimate the median difference between two genotypes and fit a 95% confidence interval around this difference.

The statistics performed associated with particular experiments is described in the figure legend for that experiment.

## Flywalker Experiments

### Hardware

The Flywalker was constructed as described in [10] with modifications. The rig consists of a frame of 80/20 supporting a sheet of 6 mm Borofloat optical glass with polished edges placed over an Andor Zyla 4.2 Magapixel sCMOS camera with an AF Nikkor 24-85mm 1:2.8-4 D lens (Nikon). On each edge of the glass were placed four Luxeon Neutral White (4100K) Rebel LED on a SinkPAD-II 10mm Square Base (230 lm @ 700mA) wired in series. Each set of lights was driven by a dedicated 700mA, Externally Dimmable, BuckPuck DC Driver (Luxeon), and all four of these drivers were connected to a single power supply. Each driver was independently adjustable.

Chambers were 3D printed by Protolabs. The ceiling of the chamber was painted with Fluon mixed with india ink, to prevent flies from walking on the ceiling. Small far-red LEDS were embedded in the walls of the chamber for Chrimson optogenetic experiments (LXM3-PD01 LUXEON). These lights were controlled by an Arduino driver that used pulsewitdth modulation to adjust light brightness. Commands were sent to this driver using a PuTTY terminal and USB serial interface. For all the experiments described here, LEDs were set at 20% brightness.

### Data Acquisition

The Flywalker was calibrated using a calibration reticle prior to use on each day. On the day of the experiment, 2-5 day old females were mouthpipetted into a clean glass tube and allowed to equilibrate for five minutes to get rid of as much dirt and food as possible to prevent contamination of the glass surface. 2-3 flies were added to the chamber by mouth pipette.

Videos were recorded using the NIS Elements AR software. A constant region of interest was defined such that the frame rate of recording was 226 fps. Each group of animals was recorded for one minute. Videos were cut to select traces where flies walked straight for >6 steps without other flies in the frame or touching the wall.

### Tracking

Flywalker videos were automatically tracked using custom software written by Imre Bartos as described in [10]. Tracking was then validated by eye and incorrect footprint calls were corrected. Summary plots were then screened by eye for gross errors and for linear traces. If traces were short (<3 traces per foot) or excessively turning, they were excluded.

### Parameters

Behavioral parameters were calculated as described in [10]. Gait parameters were defined as follows. Leg order in combination: LF RF LM RM LH RH. 1 indicates the leg is in stance phase, 0 indicates the leg is in swing phase.

### Statistics

For optogenetic Flywalker experiments, behavior was recorded for a one minute walking bout with red light illumination. Light off conditions were not possible as the white light LEDs required to generate fTIR signal contained the red wavelength used to activate our optogenetic tool. For each activation experiment, we recorded behavior from both experimental (*Trh–orotherneuromodulatoryGal4driver–⋂tsh>csChrimson*) and control (*w*^*1118*^ *⋂tsh>csChrimson*) flies. Neuromodulatory Gal4 driver lines had not been fully isogenized prior to these experiments, but two of three chromosomes (i.e., the chromosomes not containing the Gal4 itself) had been fully swapped out for those of our isogenized *w*^*1118*^ control. For each genotype, we analyzed data from flies that had been fed with ATR, the required co-factor for optogenetic activation, and flies from the same cross that had been fed on food depleted of ATR. Figures show comparisons between ATR+ and ATR-controls, as we believe these populations provide the best genetic control and had the most similar behavioral pattern. However, significant differences in parameters are consistent even when all controls are included in our multivariate model (described below).

Statistical analysis of Flywalker data was performed using custom scripts written in MATLAB and R. For each walking bout, an average was calculated for every parameter across three to five footprints per leg. For parameters that exponentially related to speed, the natural logarithm was taken of both the bout speed and parameter values. A multivariable regression model was then run on the data for every kinematic parameter. The formula for this model was as follows:

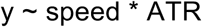

This model was designed to analyze the effects of genotype and ATR while controlling for speed, which is the largest contributor to behavioral shifts.

We also ran a version of this model that contained all control data, to validate our results:

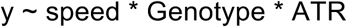

To prevent model overfitting, we selected our model based on Akaike information criterion using the R step() package.

## Functional Imaging Experiments

Functional imaging experiments on *Trh⋂tsh>opGCaMP6f, tdTomato* animals were performed and analyzed as described in [60] with the following changes.

### Analysis protocol

#### InitialimageProcessing

TIFF videos from two-photon microscopy were processed in Fiji to merge green (opGCaMP6f) and red (tdTomato) channels [106]. No brightness or contrast adjustments were performed, in order to standardize region-of-interest (ROI) selection.

#### ROISelection

The tdTomato channel was used to select ROIs containing neuronal processes, using custom Python software relying on OpenCV and Numpy libraries. Images were converted into 8-bits, color ranges were extended, and contrast was augmented to better detect ROIs. Baseline signals were subtracted and then brightness was scaled to a maximum value was 255. A blur filter was applied to the image (blur value =10), and then an Otsu Threshold was applied to binarize the grayscale image. After the image was thus thresholded, an erosion function (kernel size 5) was used to avoid the detection of overly large or small ROIs. The contours of all ROIs were detected on the eroded image and a copy of the contrast-augmented image was returned with ROI contours drawn super-imposed. A minimum threshold of 150 pixels was set on the ROI size to avoid overly small detections.

#### Fluorescenceextraction

Mean fluorescence values for the tdTomato, or opGCaMP6f channels were calculated over all ROIs combined. Baseline signals for dF/F calculations were defined as mean raw fluorescence binned over 2.5 s.

#### Synchronization

Fluorescence measurements, behavior videography, and optic flow of spherical treadmill rotations were all recorded at different frame rates. Thus, we used interpolation to upsample fluorescence signals and behavioral videography acquisition rates to that of optic flow. Optic flow and fluorescence data were then smoothed (window size 200 ms). Optic flow data was then translated into mm/s in the anterior-posterior and medial-lateral directions and into degrees/s for yaw.

#### Automatic Walking Classifier

An automatic walking classifier was used to define walking bouts. A velocity of 0.31 mm/s was empirically determined as a threshold for distinguishing between walking and standing. The minimum threshold for bout length was empirically set to 2 s.

#### Manual behavioral annotation

Videos showing a side view of the fly on the spherical treadmill were manually annotated to capture four behaviors: (1) walk, (2) stop, (3) proboscis extension reflex, and (4) groom. All frames that could not neatly be classified as one of these four behaviors were defined as (5) other.

### Statistics

#### ManuallyAnnotatedBehaviors

For each animal, the average dF/F for frames labeled a particular behavior classification was calculated. Comparisons between behaviors were made using Kruskal-Wallis testing with Dunn’s correction for multiple comparisons.

#### Timecourses

For each behavioral classification, an average time course was determined for each animal by averaging dF/F for all behavioral bouts, centering them on bout onset. Averages across all animals were then calculated, and 95% confidence intervals fit by bootstrapping.

#### Correlation Analysis

To calculate the correlation between walking velocity and dF/F, we used a Pearson correlation to calculate R.

## Acknowledgements

We thank Cesar Mendes for helping to optimize the Flywalker system, Imre Bartos for his help in updating the Flywalker analysis code, Andrew Straw for editing his motmot video collection software to make it compatible with our system, Meredith Peterson, Floris van Bruegel, Irene Kim and Michael Dickinson for assistance in building the arena hardware and data analysis approaches, Randy Bruno for assistance with analysis approaches, Laura Hermans in the Ramdya lab for her assistance with data analysis. We thank Daniel Wolpert for comments on the manuscript. This work was supported by NIH grants to R.S.M (1U01NS090514-01 and 1U19NS104655-01) the Columbia MD/PhD Training program (GM007367) and the Columbia Neuroscience Program (5T32NS064928-07). P.R. acknowledges support from the Swiss National Science Foundation (31003A_175667)

## Author contributions

C.H. and R.S.M. conceived of the project and designed the experiments; T.T. and R.H. designed and built the behavior rigs; C.H. conducted all of the experiments and performed all data analysis, with the exception of calcium imaging experiments, which were carried out by C.-L.C. and analyzed by C.H., C.-L.C., and P.R.; C.H. and R.S.M. wrote the paper and P.R. edited it.

