## Supplemental Data for "Serotonergic modulation of walking in *Drosophila*"

### Supplemental Figures

Figure S1

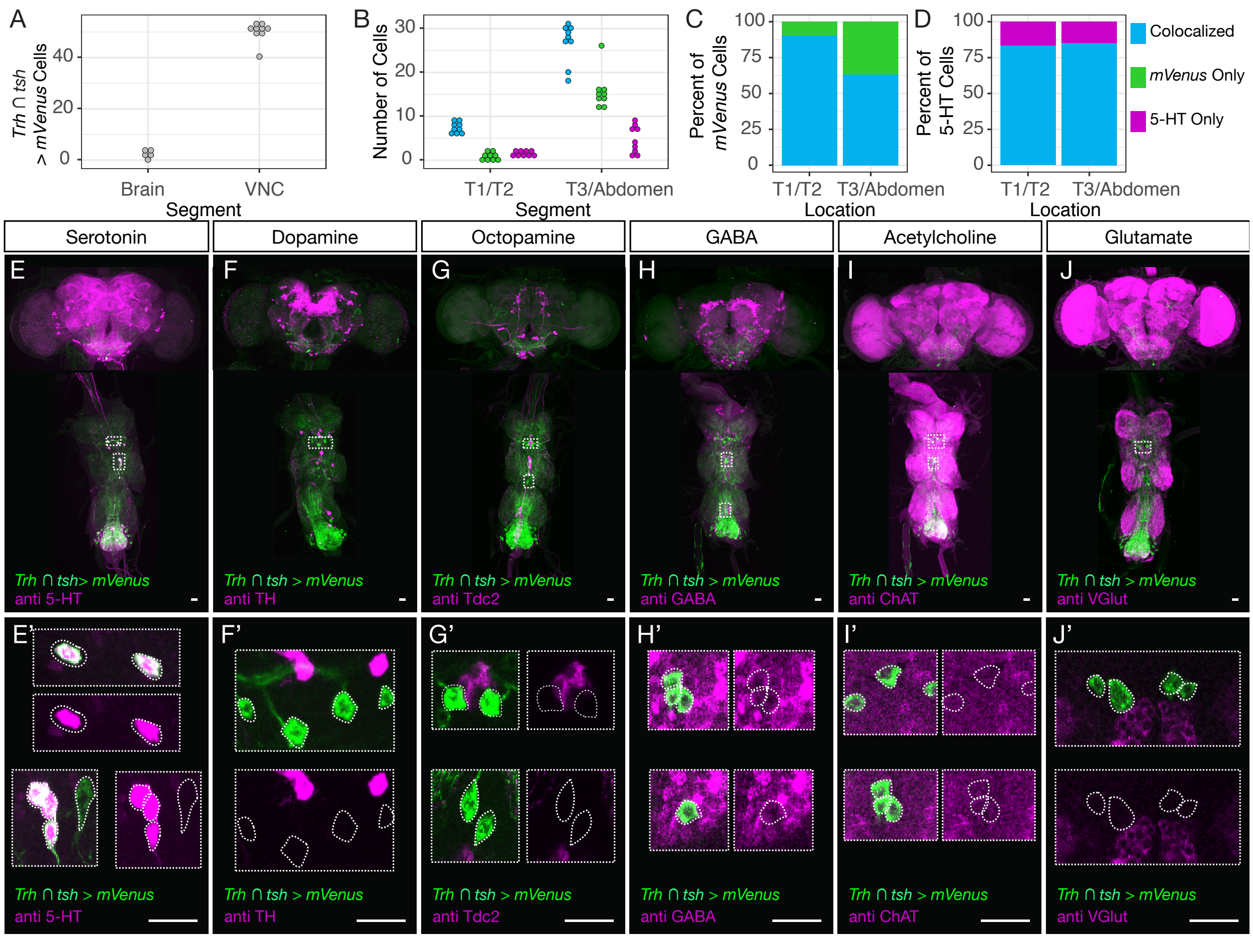


**Figure S1. Related to Figure 1. *Trh-Gal4* accurately drives expression in serotonergic VNC neurons.**

**A.** Intersection with *tsh* reliably limits *Trh-Gal4* expression to the VNC. The number of *Trh* ∩ *tsh* positive cells was quantified in the brain and the VNC (N>5). The reporter was mVenus conjugated to the UAS-csChrimson behavioral tool, and thus showed where our intersectional tool would drive expression in an activation experiment.

**B.** Quantification of the overlap between *Trh* ∩ *tsh* > *csChrimson::mVenus* and anti-serotonin (5-HT) immunostain. The number of co-localized, *mVenus* only, and 5-HT only cells was quantified separately for T1/T2 and T3/Abdomen (N=9).

**C-D.** *Trh* ∩ *tsh* has limited ectopic expression and effectively labels the majority of serotonergic neurons in the VNC. The average percent of *Trh* ∩ *tsh* > *csChrimson::mVenus* cells (C) that co-localize with 5-HT, and 5-HT positive cells that also express *mVenus* (D) is shown for the T1/T2 and T3/Abdominal regions (N=9).

**E-J.** Thoracic neurons labeled by *Trh* ∩ *tsh* > *csChrimson::mVenus* do not co-express other neurotransmitters. Maximum intensity projections of brain and VNCs of animals where the expression of *Trh-Gal4* was restricted to the VNC and co-stained for 5-HT (E), tyrosine hydroxylase (F, TH), Tyrosine decarboxylase 2 (G, Tdc2), GABA (H), Cholineacetyltransferase (I, ChAT), and the vesicular glutamate transporter (J, VGlut). White dotted boxes indicate regions that are shown in E'-J'. E'-J') Examples of *Trh* $\cap$ *tsh* > *csChrimson::mVenus* labeled thoracic neurons and neurotransmitter immunostains. Projections of 10-20 image sections show *Trh* $\cap$ *tsh* labeled neurons – dotted white outline – E') do express 5-HT, but do not express F') TH, G') Tdc2, H') GABA, I') ChAT, and J') VGlut. All scale bars represent 20 $\mu$m.

Figure S2

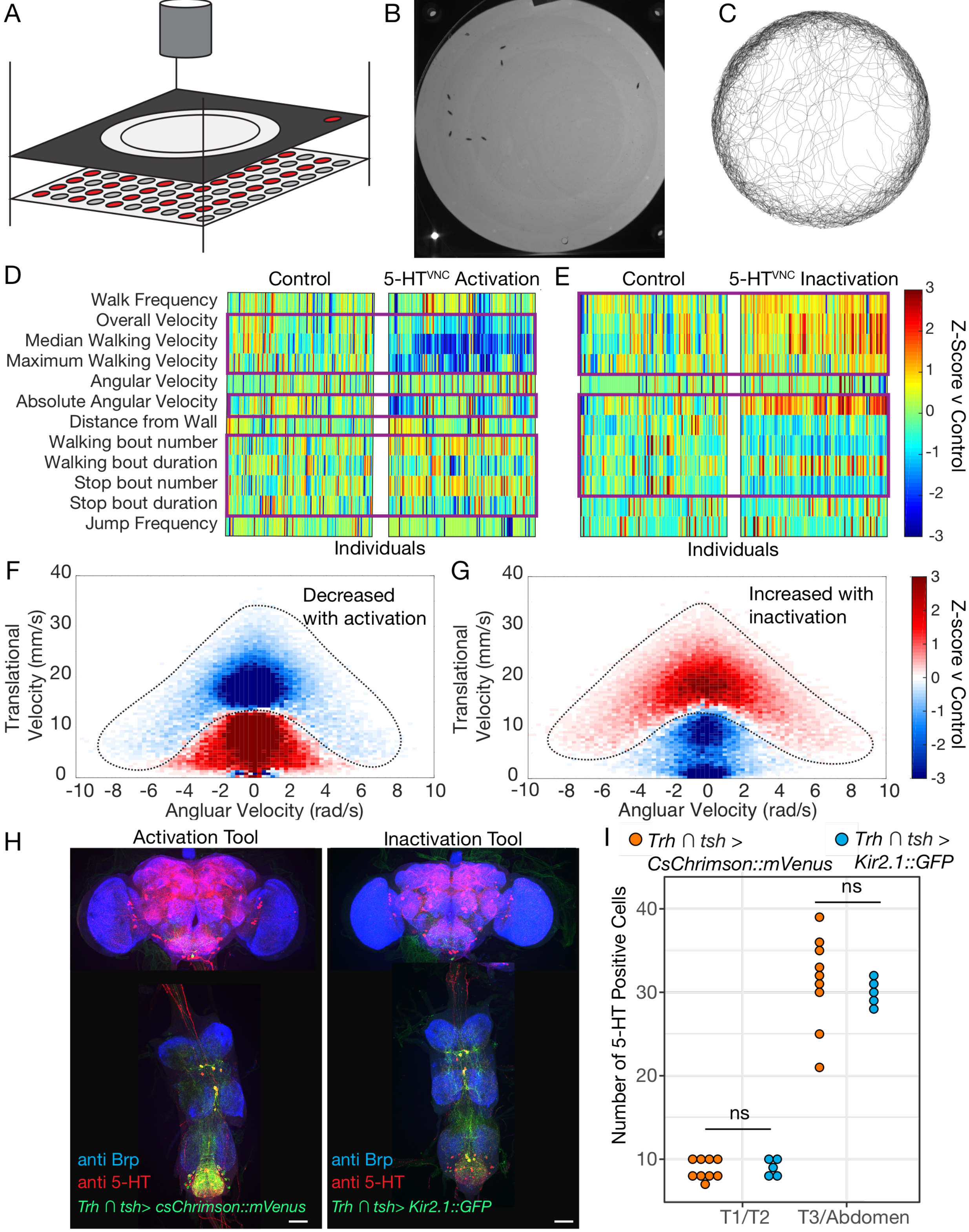


**Figure S2. Related to Figure 2. Activation and inactivation of 5-HT$^{VNC}$ neurons shifts**

**locomotor behavior.**

**A.** Schematic of the arena system used to record fly walking behavior. The system consisted of an overhead IR camera, an arena set in an aluminum plate, and a baseplate with IR backlighting and either red (for optogenetic experiments) or white LEDs.

**B.** Image of flies in the arena during a behavioral experiment.

**C.** Traces of ten animals from one five-minute recording period.

**D, E.** Heatmap of behavioral changes induced by 5-HT$^{VNC}$ neuron activation (D) and inactivation (E). Each column represents one animal, and each row one parameter. The color shows the Z-score (mean difference divided by control standard deviation) of genotype comparisons. For optogenetic experiments, behavioral change with light is compared between *Trh* $\cap$ *tsh* > *csChrimson* and background matched non-Gal4 controls (*w*$^{1118}$ $\cap$ *tsh* > *csChrimson*), both fed with all-trans-retinal (ATR). For inactivation experiments, *Trh* $\cap$ *tsh* > *Kir2.1* and *w*$^{1118}$ $\cap$ *tsh* > *Kir2.1* were directly compared. N=130 per condition for activation experiments and N=119 for inactivation experiments. Boxed are parameters where experimental animals behave significantly differently than controls p<.05. Significance calculated by Kruskal-Wallis.

**F, G.** Heatmaps showing the difference in incidence for particular velocity/angular velocity combinations with either activation or inhibition of 5-HT$^{VNC}$ neurons. F) For activation experiments, change in incidence with light was calculated for each animal and average and standard deviation was generated for each genotype (*w*$^{1118}$ $\cap$ *tsh* >*csChri*mson ATR+, *Trh* $\cap$ *tsh* >*csChrimson* ATR+). Color represents the Z-score (mean difference divided by control standard deviation) of genotype comparisons. G) Heatmaps were generated as described for activation experiments, comparing 5-HT$^{VNC}$ inactivation (*Trh* $\cap$ *tsh* > *Kir2.1)* and control (*w*$^{1118}$ $\cap$ *tsh* > *Kir2.1)* populations. N=130 per condition for activation experiments and N=119 for inactivation experiments.

**H.** Max projections of *Trh-Gal4* driving expression of fluorescent tagged activation and inhibition tools selectively in the VNC.

**I.** Quantification of the number of anti 5-HT positive cells in the T1/T2 and T3/Abdominal regions of the VNC during control (*Trh*$\cap$ *tsh* >*csChrimson::mVenus* ATR- N=9) and constitutive inhibition (*Trh* $\cap$ *tsh* >*Kir2.1::GFP* N=5) conditions. Genotypes were compared using a Welch two sample t-test. p for T1/T2 region .72, p for T3/Abdominal region .52.

Figure S3

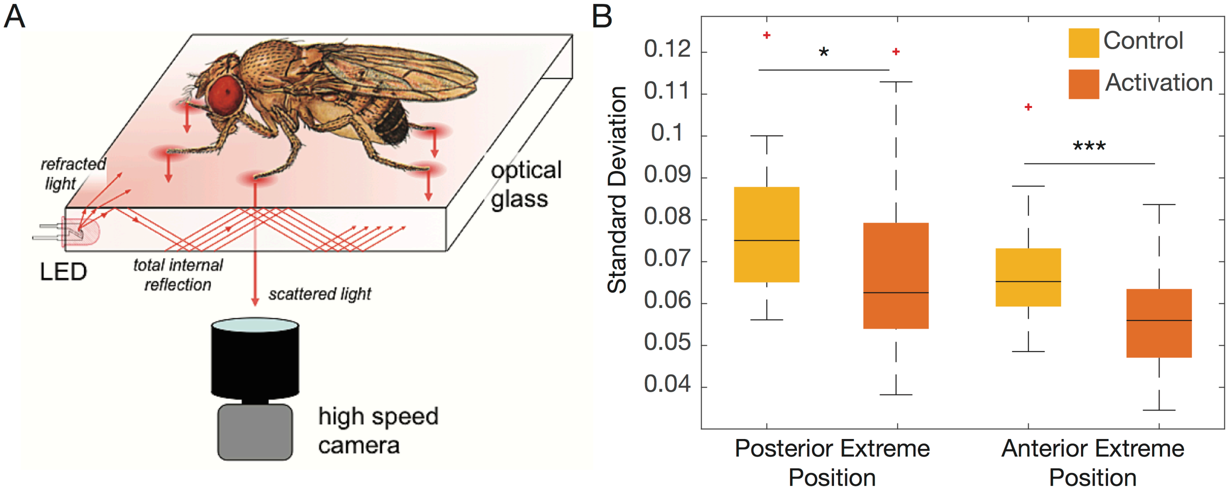


**Figure S3. Related to Figure 3. 5-HT$^{VNC}$ activation minimizes variance in foot placement.**

**A.** Schematic of the Flywalker apparatus, originally described in Mendes et al. 2013.

**B.** Boxplots showing the standard deviation of footprint position for individuals during one walking bout, either at touchdown (AEP) or lift off (PEP). N = 47 bouts from 10-23 animals for *Trh ∩ tsh >csChrimson* ATR+. N=56 bouts from 12-30 animals for *Trh ∩ tsh >csChrimson* ATR-. Genotypes were compared using a two-sample t-test. * p<.05, *** p<.001

Figure S4

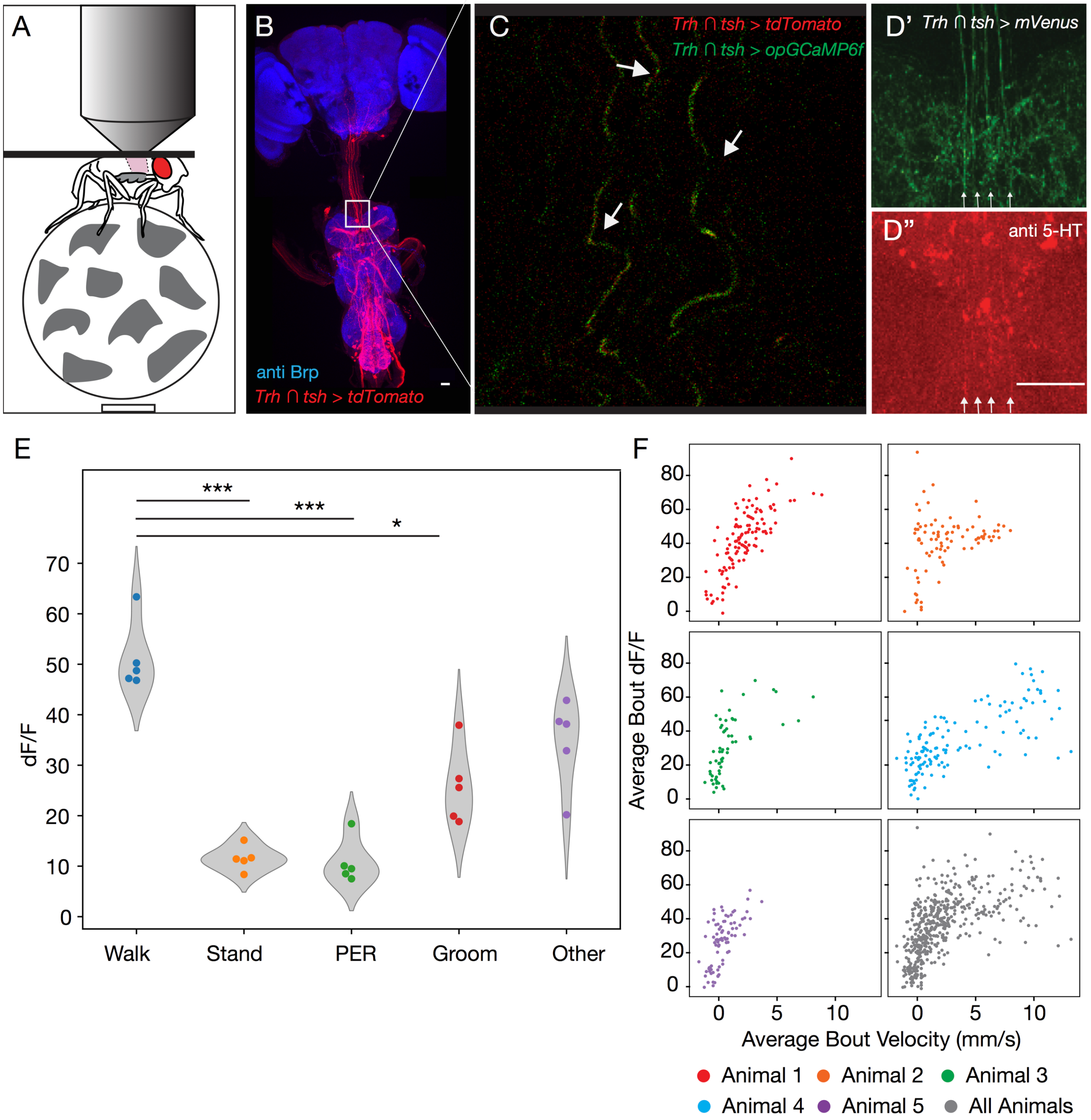


**Figure S4. Related to Figure 2. Live imaging of 5-HT$^{VNC}$ neurons during walking**

**A.** Schematic showing the set-up for VNC calcium imaging in a behaving animal. The dorsal aspect of the fly is dissected, and the fly is secured to a stage. The fly is able to freely walk on a ball suspended on an airpuff, while a two-photon microscope images fluorescence of a calcium indicator.

**B.** Expression pattern of tdTomato reporter driven by *Trh* ∩ *tsh*. This marker is expressed in the expected pattern, but has some ectopic expression in the chordotonal organ, a structure that does not express *Trh*. White box indicates the approximate window that was visualized during calcium imaging. Scale bar is 25 $\mu$m.

**C.** Single two photon image of view through recording window. Fibers - indicated by white arrows - running anterior to posterior express tdTomato (red) and codon optimized GCaMP6f (green) driven by *Trh* $\cap$ *tsh*.

**D, D'.** Image of *Trh* $\cap$ *tsh* > *csChrimson::mVenus* in the VNC shows the same fibers that we are recording from. A co-immunostain with anti 5-HT in this tissue shows that these fibers express serotonin. Scale bar = 25 $\mu$m.

**E.** The distribution of the average dF/F for each manually scored behavioral category for each animal (N=5). Significance between groups was determined by Kruskal-Wallis test with follow up Dunn's correction for multiple hypothesis testing. * p<.05, *** p<.001

**F.** Correlation between average bout velocity and dF/F for each animal, and all animals in the study (N=5). Pearson correlation coefficient for all data R=.59.

Figure S5

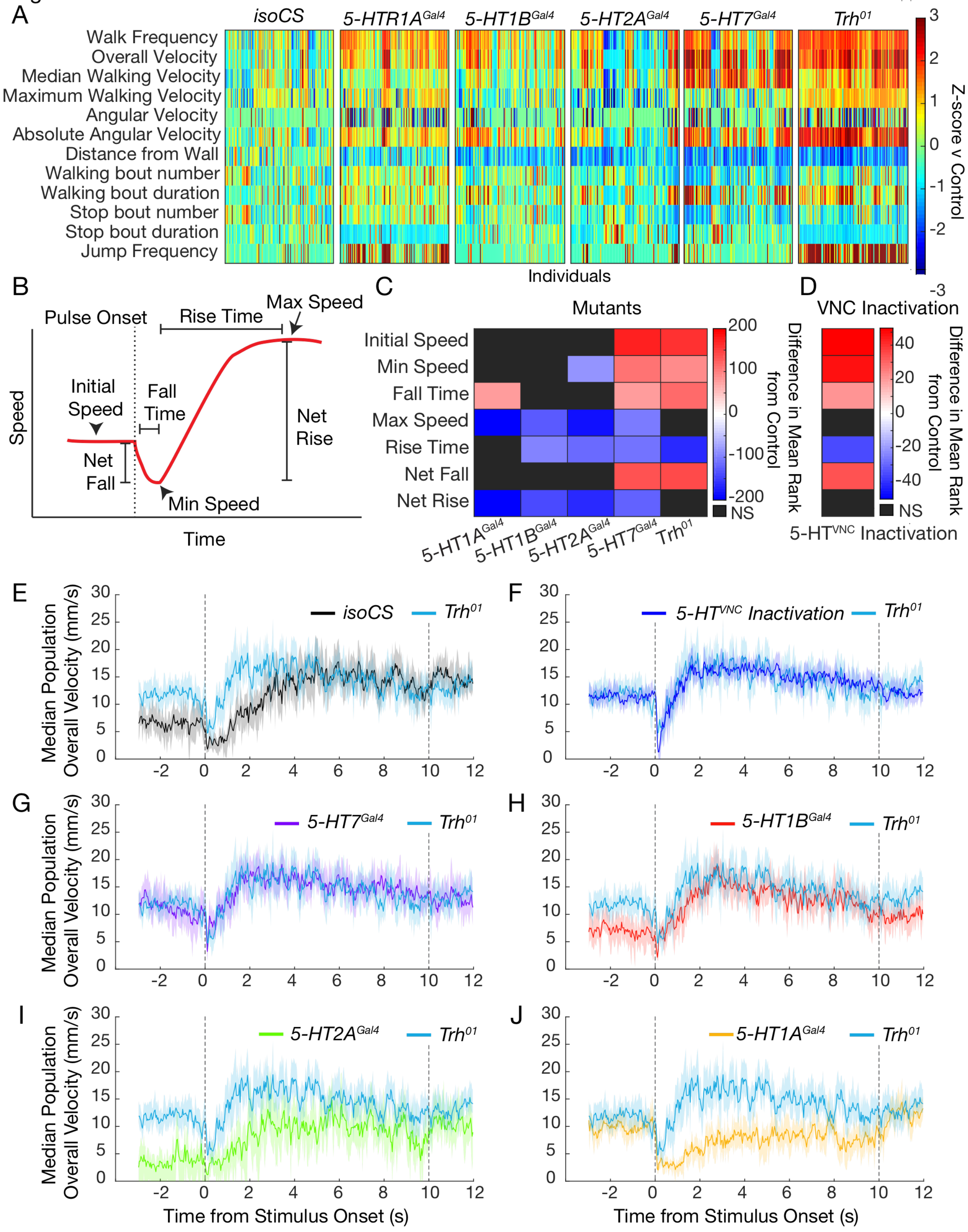


**Figure S5. Related to Figures 4 and 5. 5-HT$^{VNC}$ inactivation, mutation of *Trh*, and mutation of 5-HT receptors shifts behavioral responses to startling stimuli.**

**A.** Heatmap of behavioral differences between control animals (isoCS) and flies mutant for 5-HT receptors or *Trh* itself. Z-score for each individual was calculated using the *isoCS* group mean and sd as a control. Genotype(N) – *isoCS*(130), *5-HT1A*$^{Gal4}$ (130), *5-HT1B*$^{Gal4}$ (120), *5-HT2A*$^{Gal4}$ (100), *5-HT7*$^{Gal4}$ (120), *Trh*$^{01}$(120).

**B.** Schematic of animals' behavior in response to a sudden stimulus. This response can be divided into seven parameters that describe different phases of the response, indicated here with arrows and bounded lines.

**C.** Heatmap showing how mutation of 5-HT receptors affects the parameters depicted in panel A in response to a vibration stimulus. Genotypes were compared using Kruskal-Wallis test followed by the Dunn-Sidak correction for multiple comparisons. Plotted for every genotype is the difference in mean ranks (Kruskal-Wallis test statistics) for each parameter compared to *isoCS* control flies. Non-significant comparisons. $p>.05$, are shown in black. Genotype (N) – *isoCS*(140), *5-HT1A*$^{Gal4}$ (117), *5-HT1B*$^{Gal4}$ (130), *5-HT2A*$^{Gal4}$ (94), *5-HT7*$^{Gal4}$ (120), *Trh*$^{01}$(139).

**D.** Heatmap similar to that shown in B, except comparing inactivation of 5-HT$^{VNC}$ neurons with background matched control animals. N is 167 for *w*$^{1118}$$\cap$ *tsh >Kir2.1* and 166 for *Trh* $\cap$ *tsh >Kir2.1*.

**E-J.** Median population walking speed sampled at 30 Hz in response to vibration stimulus with 95% confidence intervals shaded. D. *Trh*$^{01}$ mutants show a blunted and shortened pause in response to novel stimulus compared to *isoCS* controls. E. The behavior of *Trh*$^{01}$ mutants replicates that caused by inactivation of 5-HT$^{VNC}$ neurons. *5-HT7*$^{Gal4}$ mutants (F, purple line) and *5-HT1B*$^{Gal4}$ mutants (G, red line) show a similar phenotype to *Trh* mutants (blue line). *5-HT2A*$^{Gal4}$ mutants (H, green line) and *5-HT1A*$^{Gal4}$ mutants (I, yellow line) have a distinct behavioral profile from *Trh*$^{01}$ mutants (blue lines).

Figure S6

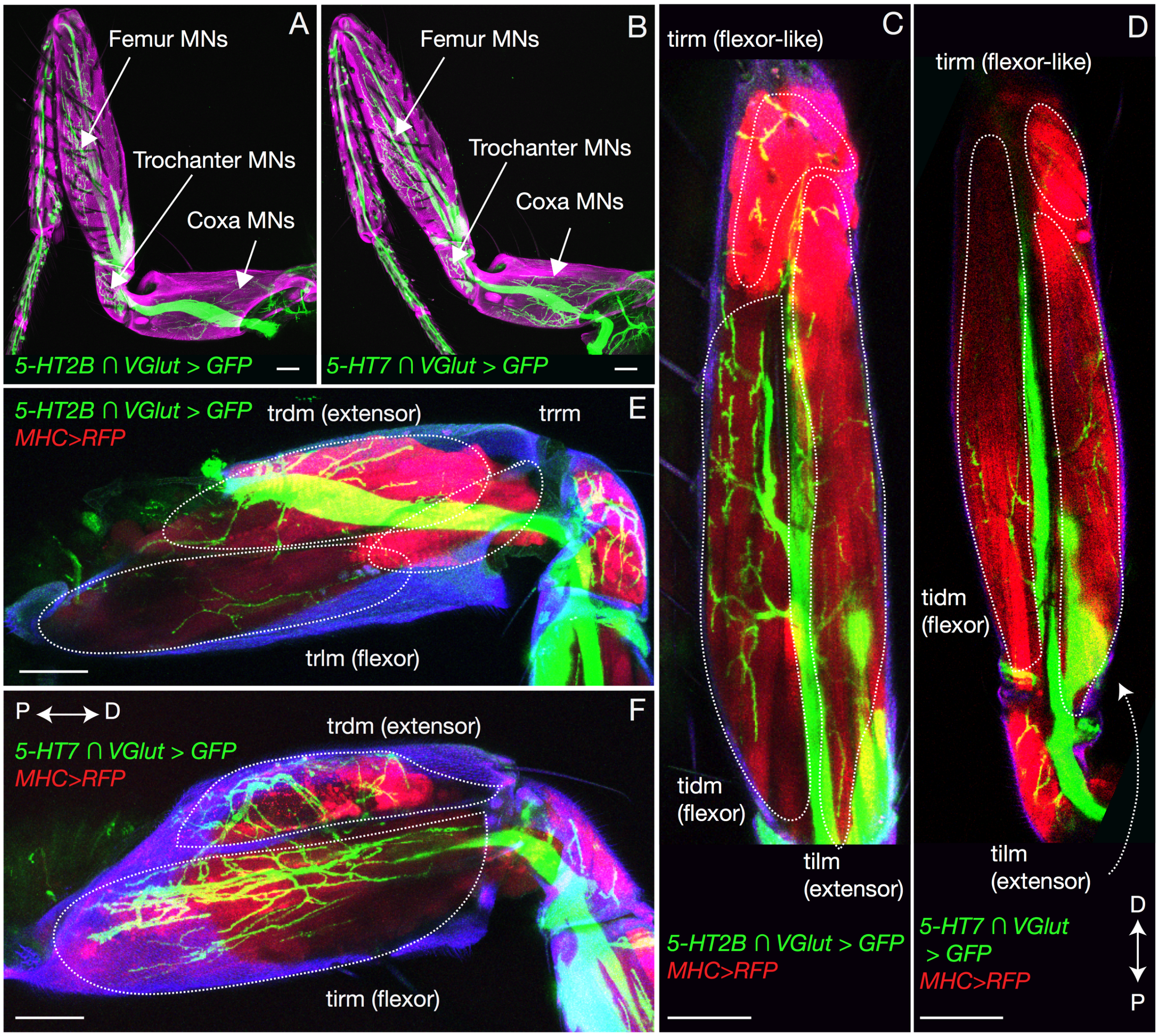


G

| Gene | Line | Protein or Gene Trap | Motor Neuron | | | | Mechanosensory | | Proprioceptive | Known Function |
|---|---|---|---|---|---|---|---|---|---|---|
| | | | Co | Tr | Fe | Ti | Prox | Dist | FeCo | |
| *5-HT1A* | MI04464 | P | | | | | | | | ↓ cAMP |
| *5-HT1B* | MI05213 | P | | | | | | | | ↓ cAMP |
| *5-HT2A* | MI00459 | P | | | | | | | | ↑ $iCa^{2+}$ |
| | MI03299 | G | | | | | | | | ↑ $iCa^{2+}$ |
| *5-HT2B* | MI05208 | P | | | | | | | | ↑ $iCa^{2+}$ |
| | MI06500 | P | | | | | | | | ↑ $iCa^{2+}$ |
| | MI07403 | G | | | | | | | | ↑ $iCa^{2+}$ |
| *5-HT7* | MI00215 | G | | | | | | | | ↑ cAMP |

**Figure S6. Related to Figure 6. 5-HT receptors are expressed in both flexor- and extensor-like muscles in the leg.**

**A and B)** Expression driven by A) *5-HT2B-Gal4* or B) *5-HT7-Gal4* after intersection with a line that limits expression primarily to motor neurons in the leg. Maximum Z-projection images show that both lines drive expression in motor neurons in the coxa, trochanter, and femur, but do not show expression in motor neurons of the tibia.  Expression in the tarsal segments is due to ectopic driver expression in some sensory populations. Scale bars represent 50 μm.

**C-F)** Motor neurons expressing 5-HT receptors innervate complementary flexor-extensor muscle pairs in the femur (C and D) and coxa (E and F). Leg muscles are labeled in red using *MHC::RFP* and are reliably identified by position and insertion points. (C and D) Max projections of multiple imaging sections through the femur shows labeled motor neuron innervation of the tibial levator muscle (tilm) – an extensor-like muscle, the tibial depressor muscle (tidm) – a flexor-like muscle, and the tibial reductor muscle (tirm) – also a flexor-like muscle. (E and F) Max projections of multiple imaging sections through the coxa shows labeled motor neuron innervation of the trochanter levator muscle (trlm) – a flexor-like muscle, the trochanter depressor muscle (trdm) – an extensor-like muscle, and the trochanter redactor muscle (trrm). Scale bars represent 50 μm. Proximal – distal orientation is indicated.

**G)** Expression in leg motor, sensory, and proprioceptive structures is annotated for all Gal4 lines tested. Strong expression is indicated as darkly colored blocks, and weaker/more selective expression is indicated by progressively lighter color. Also annotated is whether these lines were gene (G) or protein (P) traps.  Different insertion points for the same gene have highly replicable expression patterns. As described in Figure 6, receptors families with different mechanisms of action also show distinct expression profiles.
